# MISO: A Controlled Ablation of Masking, Initialization, Sampling, and Optimization for Segmentation in Volumetric Electron Microscopy

**DOI:** 10.64898/2026.06.19.733473

**Authors:** Shanmukha Vamshi Kuruba, George Stephenson, Vignesh Kasinath

## Abstract

Multi-organelle segmentation in volumetric electron microscopy (vEM) faces several challenges, including severe class imbalance, the presence of small, rare classes, and inconsistent class coverage across crops. While recent work has focused primarily on architectural design, the impact of sampling, loss functions, and masking strategies on training effectiveness remains comparatively underexplored in vEM organelle segmentation. Here, we systematically evaluate sampling strategies, loss configurations, masking approaches, and model families (CNNs and vision transformers) on the CellMap benchmark. Using 289 annotated 3D FIB-SEM crops, we establish a 32-class segmentation benchmark with stratified train, validation, and test splits, and evaluate all the methods under the same training and inference settings. Across controlled ablations, the proposed combination of repeat-factor sampling, Tversky-BCE loss, and masking achieved the strongest rare-class performance, increasing rare-class mean Dice (mDice) from 0.3244 under uniform sampling to 0.3409. This corresponds to an absolute gain of +0.0165 mDice and a 5.1% relative improvement, while preserving comparable performance on common classes. Overall, we find that sampling, loss design, and masking contribute as much to performance variation as the choice of architecture, highlighting the importance of training-recipe design alongside model architecture in vEM organelle segmentation.

## 1. Introduction

Volumetric electron microscopy (vEM) enables imaging of cellular structures at nanometer resolution Peddie, Genoud, Kreshuk, Meechan, Micheva, Narayan, Pape, Parton, Schieber, Schwab et al. (2022). One such modality of vEM is Focused ion beam scanning electron microscopy (FIB-SEM) Xu, Hayworth, Lu, Grob, Hassan, García-Cerdán, Niyogi, Nogales, Weinberg and Hess (2017), which now produces multi-terabyte volumes that capture entire cells and tissues, allowing large-scale analysis of cellular structures. However, extracting biological information from these volumes requires dense semantic segmentation of numerous organelles and membrane-bound structures. As shown in Fig 1, this task is particularly challenging because of the shallow contrast profile of the different object classes in raw EM images, which in addition exhibit highly overlapping intensity profiles, making them difficult to distinguish using pixel intensity alone. Consequently, segmentation of dense EM volumes remains largely manual or semi-automated and represents a major bottleneck in the analysis pipeline. Reliable automation would enable high-throughput phenotypic studies, mapping of organelle interactions, and large-scale comparisons across biological conditions.

**Fig. 1:**
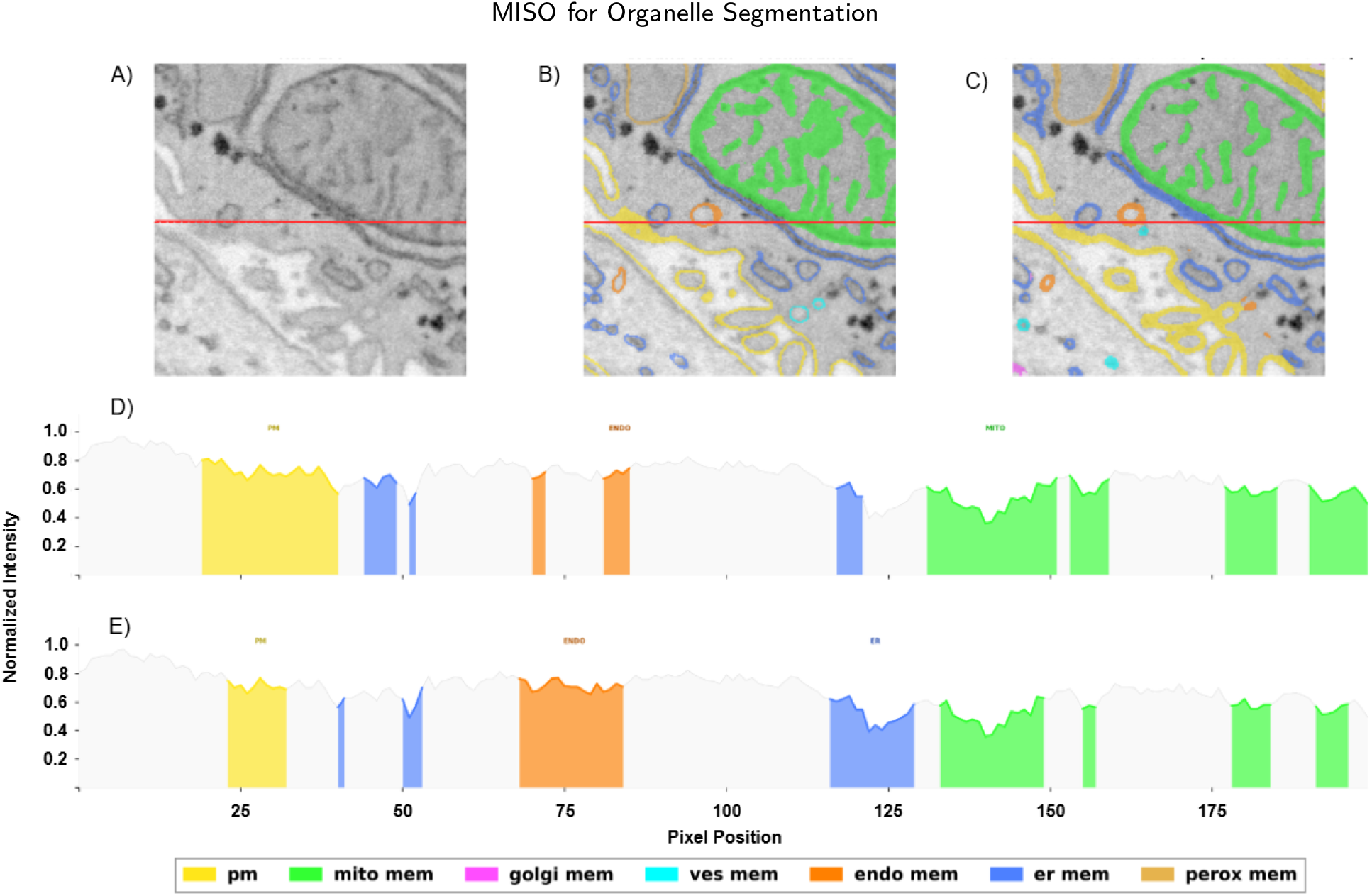
A) Raw EM image, B) ground truth annotations, and C) model predictions for seven membrane classes in a liver tissue sample. D) and E) Intensity profiles Ground truth and prediction respectively, extracted along the dashed line reveal that different membrane types exhibit highly overlapping intensity distributions, making them indistinguishable by pixel values alone.

Deep learning has driven significant progress in the automated segmentation of cellular structures. Some of the early vEM segmentation methods were based on U-Net architectures, which have since been expanded to automated pipelines such as nnU-Net, context-aware CNNs such as DeepLabV3+, and transformer-based architectures designed to capture long-range spatial dependencies Ronneberger, Fischer and Brox (2015); Isensee, Jaeger, Kohl, Petersen and Maier-Hein (2021); Sreekumar, Palanisamy and Swaminathan (2023); Zhang, Lin, Wang, Chu, Yang, Xiao, Lin and Liu (2024). Recent improvements in data collection and annotation, such as the richly annotated OpenOrganelle Heinrich, Bennett, Ackerman, Park, Bogovic, Eckstein, Petruncio, Clements, Pang, Xu et al. (2021), along with benchmarks like MitoEM Wei, Lin, Franco-Barranco, Wendt, Liu, Yin, Huang, Gupta, Jang, Wang et al. (2020) and CellMap CellMap Consortium (2024), have enabled systematic evaluation of vEM Segmentation methods. However, research in this area has focused largely on architectural innovation, while the influence of training configuration and supervision strategy remains less explored.

Such an approach, however, overlooks several key properties of vEM segmentation. First, class coverage is highly uneven. Each crop (a fixed-size 3D sub-volume extracted from the raw electron microscopy volume) is fully annotated, but only for the organelles it contains. It is rare for a single crop to contain instances of all organelle classes. Common classes such as the endoplasmic reticulum are present in many crops, whereas others, such as Golgi membranes, appear in very few. Models must therefore handle both missing classes within crops and severe global imbalance, where rare classes contribute less than 0.32% of voxels. Second, the benchmark definition itself affects results. Choices such as which classes to include, how to split the data, and which metrics to report can change model rankings without changing the model. A method that ranks first on mean Dice may rank much lower on rare classes, making comparisons ambiguous without clear evaluation criteria. Third, the training recipe strongly affects performance. Sampling strategy, loss function, and masking all interact with class imbalance and uneven class coverage. For example, changes in sampling strategy alone can alter rare-class Dice by more than 6 percentage points, which raises two key questions: How much does the training recipe matter compared to architecture? And what combination of sampling, loss, and masking best handles this setting?

We address these questions through controlled experiments across four axes: (i) sampling strategy, (ii) loss design, (iii) masking strategy, and (iv) architecture choice (with and without pretraining). Each axis is optimized sequentially on the validation set while keeping earlier choices fixed, thereby avoiding test-set leakage. We find that training-recipe choices contribute as much to performance variation as architecture. Differences between models are most pronounced in rare-class segmentation, where convolutional models benefit from their local inductive bias compared to transformer-based approaches.

Our contributions are as follows. We introduce a benchmark with stratified train, validation, and test splits, audited for leakage and distributional balance. We present a controlled evaluation across sampling, loss design, masking, and architecture choices for imbalanced EM segmentation. We believe the proposed training configuration, combining context-preserving repeat-factor sampling, recall-biased supervision, and pixel-wise uncertainty-guided masking, can be incorporated into other segmentation architectures as a general strategy for improving rare-class performance. Finally, we provide practical guidelines for training under severe class imbalance, including when pretraining helps and which losses are most effective for rare classes.

The rest of the paper is organized as follows. Section 2 reviews prior work. Sections 3 describe the dataset, experimental setup and defines the Ablations. Section 4 presents results, and Section 5 discusses implications and limitations of the study, and Section 6 concludes the paper.

## 2. Related Work

### Volumetric EM Segmentation

FIB-SEM imaging has enabled high-resolution reconstruction of cellular structures, making it well suited for 3D organelle analysis Xu et al. (2017). Within this setting, the COSEM project extended segmentation to multiple classes by using 3D U-Nets trained to predict signed tanh boundary distances Heinrich et al. (2021). Previous work on EM benchmarks was narrower in scope, focusing on specific tasks such as membrane segmentation in ISBI 2012 Ciresan, Giusti, Gambardella and Schmidhuber (2012) or synaptic connectivity in CREMI Buhmann, Sheridan, Malin-Mayor, Schlegel, Gerhard, Kazimiers, Krause, Nguyen, Heinrich, Lee et al. (2021). More recent efforts, such as MitoEM, have scaled evaluation to mitochondrial instance segmentation Wei et al. (2020). Building on these task-specific benchmarks, the CellMap challenge introduces a broader multi-class setting with 289 annotated crops from 22 datasets and evaluation across 48 (atomic and composite) classes CellMap Consortium (2024).

### Architectural Evolution

Full 3D architectures offer strong representational capacity for volumetric EM segmentation, but their memory and computational costs can limit scalability. As a result, 2D and 2.5D approaches remain widely used when computational efficiency and large-scale experimentation are important Zhang, Liao, Ding and Zhang (2022). The original U-Net remains a standard architecture for biomedical segmentation Ronneberger et al. (2015), while extensions such as U-Net++ and nnU-Net improve skip connections and training configuration Zhou, Siddiquee, Tajbakhsh and Liang (2019); Isensee et al. (2021). Transformer-based models, including SwinCell Zhang et al. (2024) and UNETR Hatamizadeh, Tang, Nath, Yang, Myronenko, Landman, Roth and Xu (2022), introduce global context through attention mechanisms. However, EM images differ substantially from natural images, and many target classes are both tiny and sparsely represented. As a result, it remains unclear whether pretrained encoders or transformer-based models consistently outperform well-optimized convolutional baselines in multi organelle segmentation setting.

### Sampling and Class Imbalance

Severe class imbalance is a central challenge in organelle segmentation. Many biologically important classes occupy only a small fraction of the volume. In dense prediction tasks, class imbalance occurs both within individual crops, where rare classes occupy very few voxels, and across crops, where many samples contain little or no annotation for those classes. Under uniform sampling, many patches contain little useful signal for rare classes. Common approaches address this by biasing sample selection, for example, through weighted sampling Conrad and Narayan (2023), online hard patch mining He, Zhou, Zhou and Chen (2021), or class-balanced mini-batches Shen, Lin and Huang (2016). Foreground-guided methods ensure exposure by centering patches on annotated classes. Class-balanced batching enforces representation across rare, medium, and common classes. Repeat-factor sampling (RFS), introduced for LVIS (a large-vocabulary instance segmentation benchmark with long-tailed category distributions), increases the sampling frequency of images containing rare categories using a frequency-dependent repeat factor Gupta, Dollar and Girshick (2019). These approaches reflect a common trade-off between increasing exposure to rare classes and preserving spatial diversity and context.

### Loss Functions

Extreme class imbalance makes loss design critical. Dice loss Milletari, Navab and Ahmadi (2016) directly optimizes pixel-level overlap between predicted and ground truth segmentation masks, and is less sensitive to class frequency. Focal loss Lin, Goyal, Girshick, He and Dollár (2017) addresses class imbalance by down-weighting predictions the model makes with high confidence and focusing training on those where the model is uncertain or wrong. In applications where missed target structures are particularly undesirable, recall-biased objectives are often preferred. The Tversky loss Salehi, Erdogmus and Gholipour (2017) extends Dice by introducing separate weights for false positives and false negatives, allowing explicit control over this trade-off. Recent work combines overlap-based and pixel-wise losses to improve both stability and performance Isensee et al. (2021).

### Masking Strategies

Masking strategies modify supervision to improve training under noise and imbalance. At the input level, masked supervised learning introduces reconstruction objectives, where randomly masked pixels force the model to use broader spatial context Zunair and Hamza (2022). At the annotation level, box-based masking restricts supervision to regions around labeled classes, reducing unintended correlations with background Song, Huang, Ouyang and Wang (2019). Previous SSL approaches often use confidence or uncertainty thresholds to filter unreliable pseudo-labels Ghazal, Tanha, Shahi and Roshan (2025), which can stabilize training but may also discard moderately uncertain samples that remain informative. Ghazal et al. (2025) address this limitation using dynamic entropy masking and entropy-based weighting, filtering only extremely uncertain pseudo-labels while retaining informative uncertain samples.

## 3. Materials and Methods

### 3.1. CellMap Benchmark and Dataset Audit

For this study, we use the CellMap organelle segmentation dataset from the Janelia CellMap Challenge CellMap Consortium (2024). The dataset contains 289 fully annotated 3D FIB-SEM crops from 22 volumes, covering 35 atomic organelle classes. These include major classes such as mitochondria, the ER, the Golgi apparatus, vesicles, lysosomes, and the nucleus, each with a membrane and lumen component. Annotations are provided at the crop level rather than as dense patches, so each sample corresponds to a distinct spatial region with varying class presence. Following the CellMap evaluation protocol, we construct crop-level training and evaluation splits.

#### Train/Validation/Test Split

To create the benchmark, we split the 289 crops from 22 volumes into 214 training crops (74.0%), 38 validation crops (13.2%), and 37 test crops(12.8%). We stratify the split at the volume level to preserve the crop distribution within each source volume. For volumes containing multiple crops, we use an approximate 75/12.5/12.5 train/validation/test split. A schematic example of the resulting crop distribution within a volume is shown in Fig 2. For volumes containing a single crop, we assign the crop entirely to either the validation or test set. A detailed split of the crops used for training and evaluation is provided in Supplemental Table 1.

**Fig. 2:**
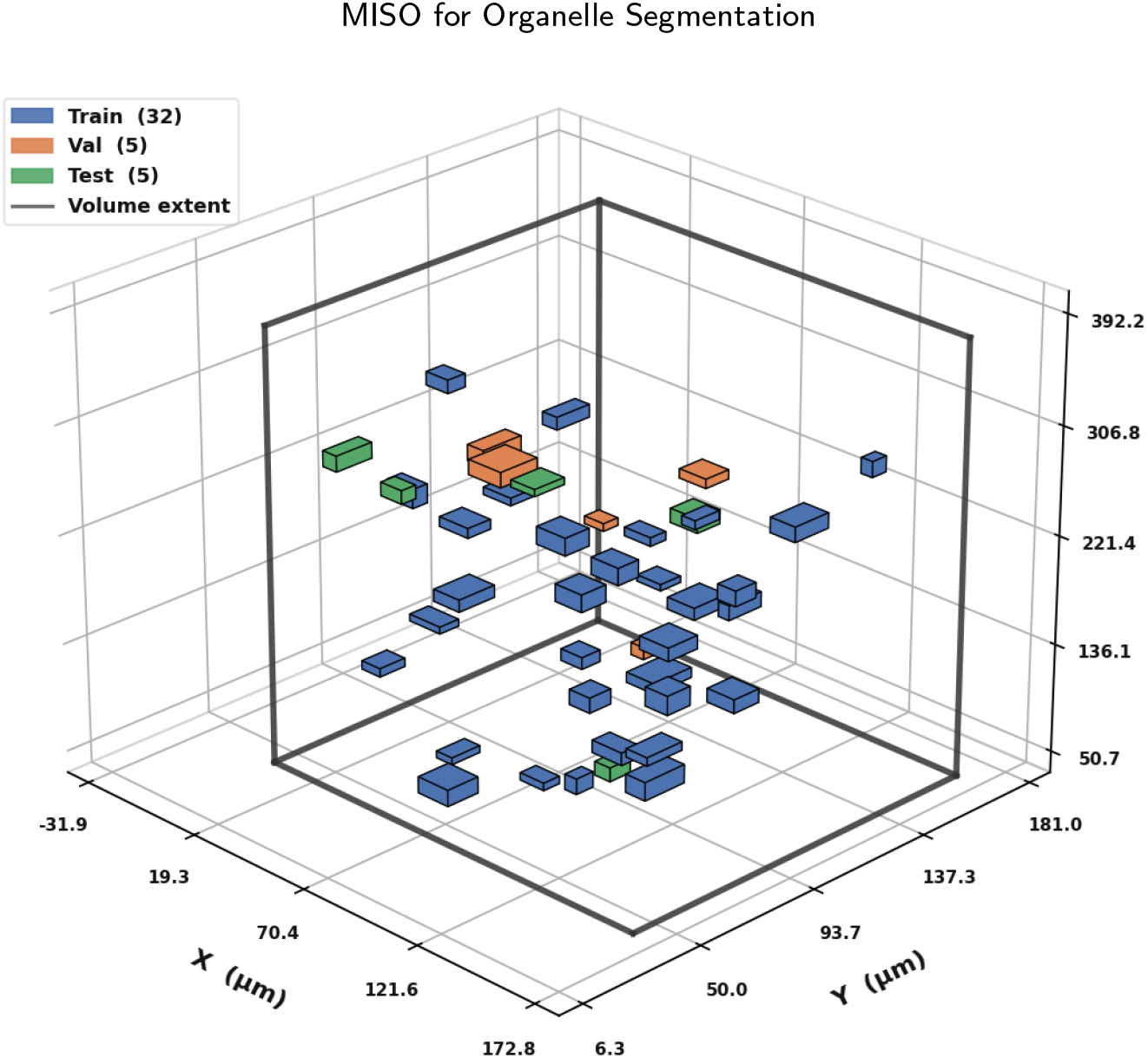
Schematic overview of crop distribution within a representative 3D volume. Boxes mark the approximate locations of annotated crops and are colored by data split. The wireframe outlines the full volume.

#### Class Coverage and Evaluation Exclusions

A class-level analysis showed that nuclear heterochromatin (nhchrom), nuclear euchromatin (nechrom), and nucleolus (nucleo) appear in only two crops each. These classes cannot be reliably distributed across train, validation, and test splits while preserving volume-level stratification. Their exclusion from training is therefore due to limited dataset coverage rather than the split design. We therefore exclude these classes from both training and evaluation, resulting in a 32-class benchmark (Table 1). Since these classes are sub-components of the nucleus, their removal does not affect higher-level structural context.

**Table 1.** The dataset contains 35 atomic class labels. Of these, 32 classes are used for training and evaluated independently in the benchmark. Classes marked with ^†^ appear in ≤ 2 crops and are excluded from independent evaluation.

| *Background & organelles (I)* |  | *Organelles (II)* |  | *Nucleus & cytoskeleton* |  |
| --- | --- | --- | --- | --- | --- |
| ecs | extracellular space | lyso_mem | lysosome membrane | ne_mem | NE membrane |
| pm | plasma membrane | lyso_lum | lysosome lumen | ne_lum | NE lumen |
| cyto | cytoplasm | ld_mem | lipid droplet mem. | np_out | NP outer ring |
| mito_mem | mito. membrane | ld_lum | lipid droplet lumen | np_in | NP inner ring |
| mito_lum | mito. lumen | er_mem | ER membrane | hchrom | heterochromatin |
| mito_ribo | mito. ribosomes | er_lum | ER lumen | echrom | euchromatin |
| golgi_mem | Golgi membrane | eres_mem | ER exit site mem. | nucpl | nucleoplasm |
| golgi_lum | Golgi lumen | eres_lum | ER exit site lumen | *Excluded*<sup>†</sup> |  |
| ves_mem | vesicle membrane | mt_out | microtubule outer | nhchrom<sup>†</sup> | nucl. heterochromatin |
| ves_lum | vesicle lumen | mt_in | microtubule lumen | nechrom<sup>†</sup> | nucl. euchromatin |
| endo_mem | endosome membrane | perox_mem | peroxisome mem. | nucleo<sup>†</sup> | nucleolus |
| endo_lum | endosome lumen | perox_lum | peroxisome lumen |  |  |

#### Split Validation

The train, validation, and test sets share no crops, and no original validation samples are used for training. Distributional alignment is confirmed using two tests on the 32-class annotation profiles. Train-test alignment is supported by both distributional tests on the 32-class annotation profiles. The Jensen-Shannon divergence is low (0.0076), and a 500-permutation MMD test gives an unbiased estimate of MMD^2^ = -0.00162 (p = 0.666), consistent with no detectable train-test discrepancy. We observe a train-validation shift (9.8 vs. 7.2 classes per crop, Wasserstein distance 2.58), driven mainly by annotation density rather than class composition. This reflects the original dataset and does not affect train-test alignment.

#### Class Distribution

The 32 retained classes span four orders of magnitude in voxel count (Fig 3). Large classes such as cyto (38.00%), ecs (18.86%), and nuc (17.76%) dominate the dataset, while 18 rare classes fall below 0.32% of voxels, down to eres_lum (0.026%), np_in (0.014%), and mito_ribo (0.008%). In this dataset, class imbalance arises in two ways. Some classes are rare across the dataset, while others appear in many crops but occupy only a small fraction of each crop.

**Fig. 3:**
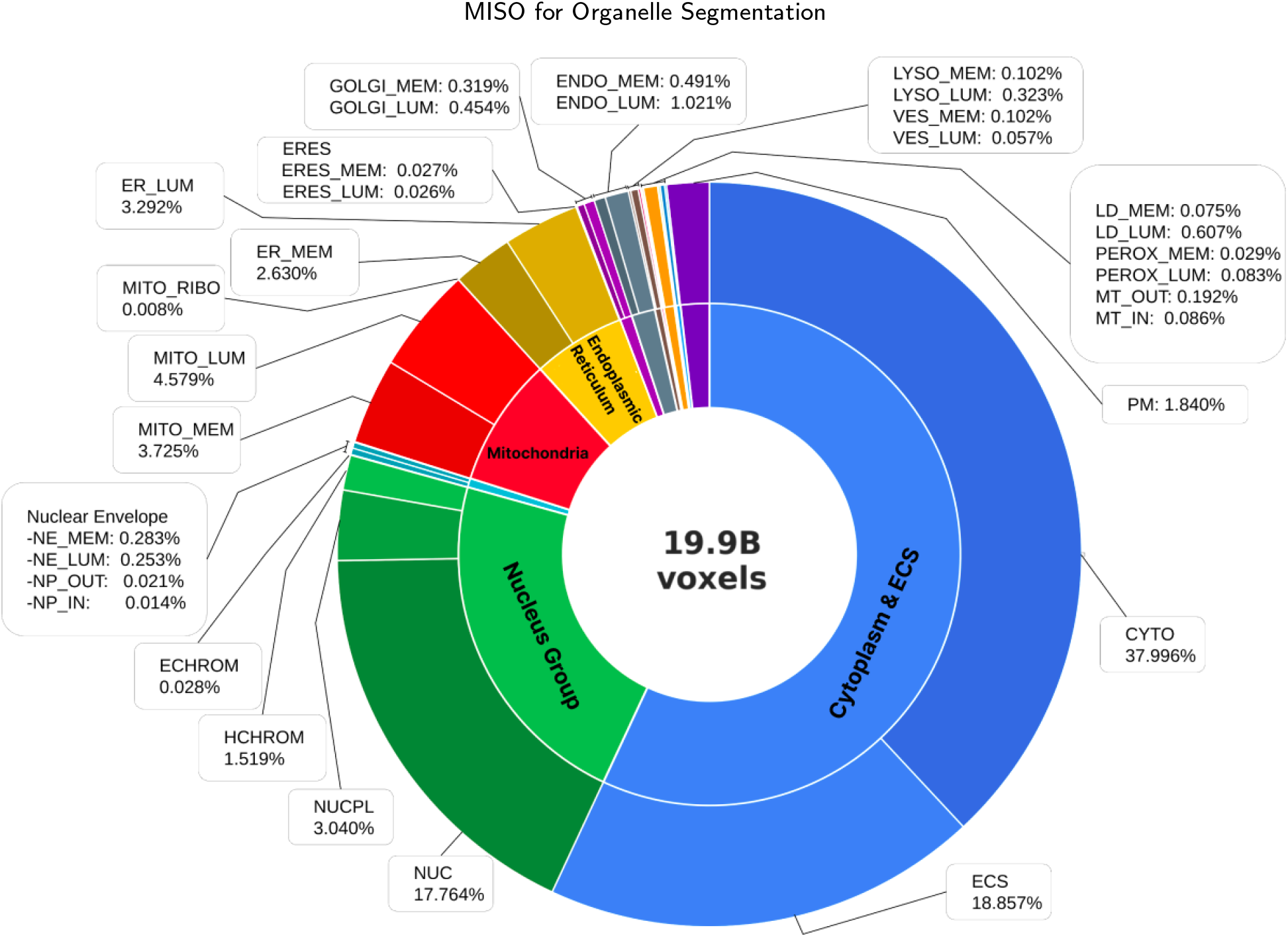
Distribution of foreground voxels across classes in the training crops.

Our final benchmark evaluates 32-class segmentation over 289 crops (214 train, 38 val, 37 test) from 22 volumes. We report overall mean Dice, along with common- and rare-class Dice, to ensure performance is not dominated by large classes.

### 3.2. Experimental Setup

#### 2D Experimental Design

We use 2D slice-based segmentation for all experiments instead of fully 3D models. Although 3D models can use inter-slice context, they require much more memory and computation Brügger, Baumgartner and Konukoglu (2019), making large controlled ablations across sampling, loss, masking, and architecture settings difficult under fixed compute resources. Since the goal of this study is to compare training configuration components in a controlled setting, all experiments use the same 2D setup. Our focus is therefore on understanding how sampling, loss design, masking, and architecture affect segmentation under extreme class imbalance, rather than maximizing absolute benchmark performance using large volumetric models.

#### Implementation Details

Each model processes single-channel 256 × 256 patches, with each patch corresponding to a 2D slice from a FIB-SEM volume. During training, each patch is augmented with random flips and rotations up to 90_°_. For crops containing rare classes, we additionally apply elastic deformation and intensity jitter with probability 0.7 to reduce overfitting on the limited rare-classes. We train all models using the AdamW optimizer Loshchilov and Hutter (2019) with an effective batch size of 64, achieved through gradient accumulation. We use automatic mixed-precision training with PyTorch AMP, which performs selected CUDA operations in lower precision to reduce memory usage and improve training efficiency. Models trained from scratch use random initialization, a learning rate of 10^−4^, weight decay of 10^−4^, and a OneCycleLR schedule Smith and Topin (2019) with 5% warmup. For pretrained models, we freeze the early encoder stages and use layer-wise learning-rate decay with a decay factor of 0.75 per depth, together with a 10% warmup schedule. We run all ablation experiments for 60 epochs, and each model is trained on a single NVIDIA RTX 3090 GPU with 24 GB of memory.

#### Evaluation Metrics

We measure model performance using mean Dice across the 32 retained classes, with mean IoU (Intersection over Union), precision, and recall as supporting metrics. We also report results separately for rare (18 classes with < 0.32% voxels, e.g., `eres_mem, eres_lum, golgi_lum`) and common classes (remaining 14 classes) Maier-Hein, Reinke, Godau, Tizabi, Buettner, Christodoulou, Glocker, Isensee, Kleesiek, Kozubek et al. (2024). We use deterministic tiling with *z*-stride 4 to balance cost and coverage for Inference. To show when aggregate metrics obscure rare-class behavior, we report both overall and rare-class Dice.

### 3.3. Ablation Study Design

Our study is organized around four experimental axes, with each stage modifying a single design component while eeping the best-performing configuration from the previous stage fixed. The sequential setup allows for reduced computational cost compared to a full grid search while allowing the effect of each component to be evaluated independently. Across all experiments, we keep the effective batch size fixed at 64. Sampling, loss, masking, and architecture experiments use a batch size of 16 with 4 gradient accumulation steps. All ablations are evaluated on the validation set using mean Dice, reported overall and stratified by class frequency. The overall training pipeline used across these experimental axes is shown in Fig 4.

**Fig. 4:**
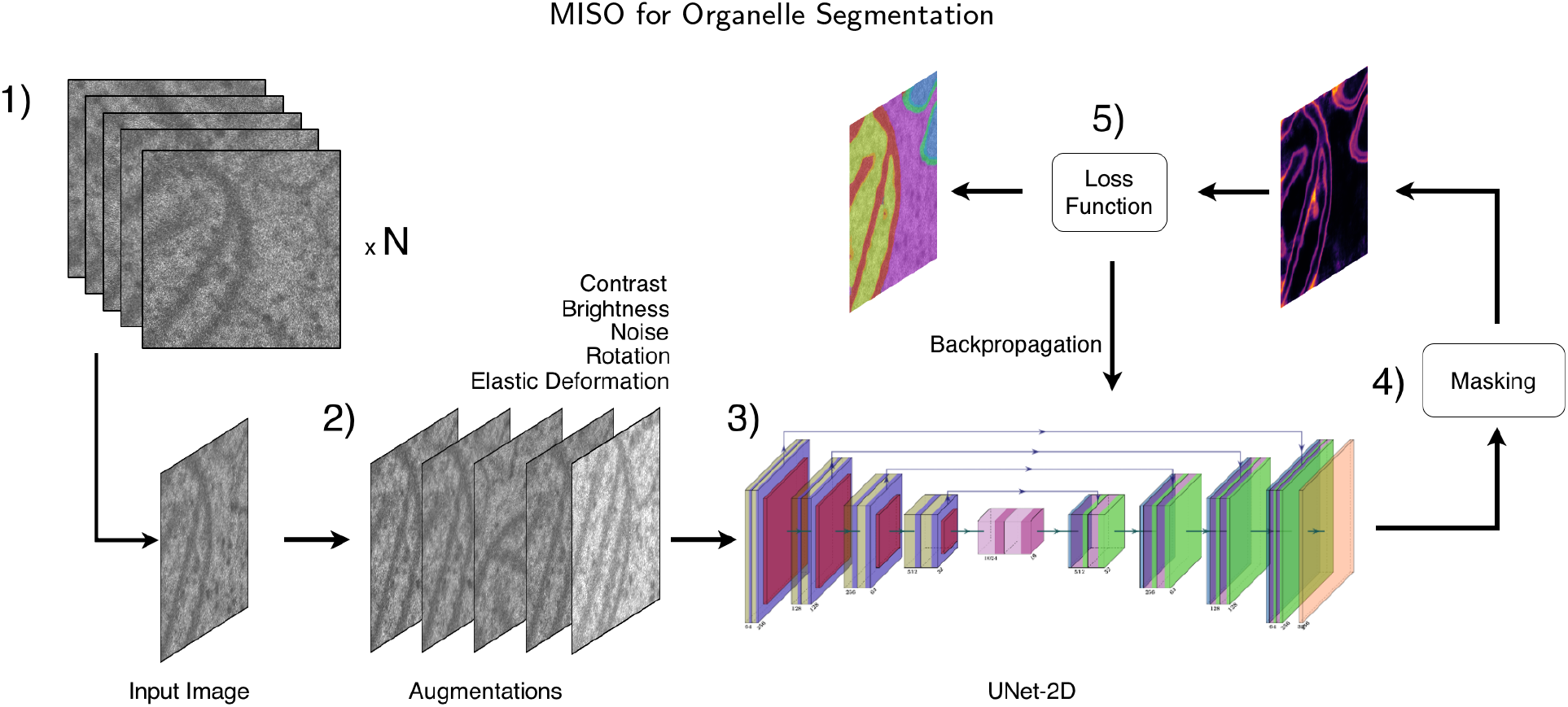
Complete training pipeline for multi-organelle segmentation from FIB-SEM images. (1) Each 3D crop is assigned a repeat factor according to the sampling strategy, increasing the probability of selecting crops that contain rare or underrepresented classes. During training, a crop is sampled from this weighted pool, after which a 2D slice is extracted from the selected crop. (2) Data augmentations including contrast adjustment, brightness variation, noise injection, rotation, and elastic deformation are carried out of the extracted 2D slices. (3) The augmented slices are processed by a 2D UNet encoder-decoder architecture. (4) Pixel-wise masking is applied before loss computation. (5) The masked predictions are used for loss computation and model optimization.

#### Sampling Strategy Ablation

The CellMap dataset presents extreme class imbalance as its central training challenge. The 32 training classes span a ∼4750:1 ratio by voxel count (e.g., cyto = 38%, `mito_ribo` = 0.008%, `np_in` = 0.014%). We apply all sampling strategies at the crop and z-slice level. A *crop* is a 3D volume, from which 256 × 256 2D patches are extracted at sampled z-slices. At each training iteration, we first select a 3D crop according to the sampling strategy. We then sample a z-slice from that crop and extract a 256 × 256 2D patch. For crop-level strategies, we change only the crop-selection probability. The z-slice and spatial location are sampled uniformly. For foreground-guided strategies, we center each patch near annotated foreground pixels. All experiments use 2D UNet, Dice loss, no masking, and 60 epochs with 2,000 iterations per epoch.

- **Uniform random sampling**. We select crops uniformly at random. Each crop therefore has the same sampling probability, regardless of its class content.
- **Class-aware crop sampling**. Here we up-weight crops that contain rare classes. We use a 70/30 mixture of class-aware and uniform sampling probabilities. Within a selected crop, z-slice and spatial location are sampled uniformly.

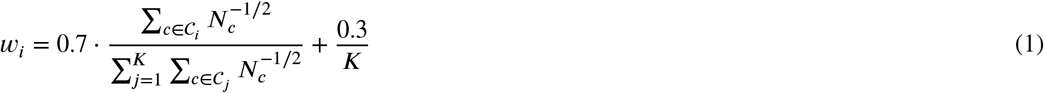

where *C*_*i*_ is the set of annotated classes in crop *i, N*_*c*_ is the global voxel count of class *c, K* is the number of crops, and Σ_*i*_ *w*_*i*_ = 1.
- **Repeat-factor sampling**. We assign each crop a repeat factor based on the frequency of its rarest annotated class Gupta et al. (2019). We adapt the LVIS repeat-factor sampling approach by using the minimum class frequency per crop, allowing the rarest class in each crop to determine the sampling rate.

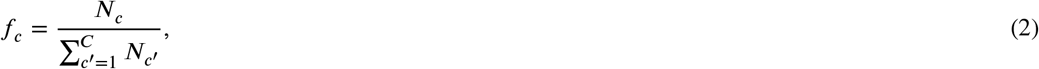

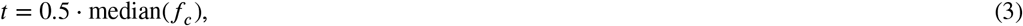

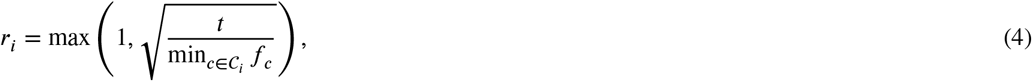

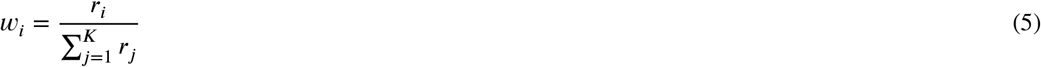

where *f*_*c*_ is the global frequency of class *c, t* is the threshold, *r*_*i*_ is the repeat factor for crop *i*, and Σ_*i*_ *w*_*i*_ = 1.
- **Foreground-guided patch mining**. We would like to note here that crop-level weighting does not guarantee that sampled patches contain rare-class pixels. To address this discrepancy, we use foreground-guided patch mining to pre-index all foreground locations by scanning label volumes and storing (crop, *z, y, x*) coordinates for each class. During training, a class is sampled with inverse-frequency weighting. We then select the foreground locations uniformly, and extract a patch centered at that point with a spatial jitter of ±32 pixels
- **Class-balanced batch**. Inspired by class-balanced mini-batch sampling approaches Shen et al. (2016), we sort classes by voxel count and divide them into three tiers: rare, medium, and common. Each batch contains 25% samples from each tier and 25% from standard random sampling, enforced deterministically by cycling through tiers. Within each tier, we sample classes using 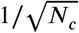 weighting.
- **Hybrid**. We pick each sample with probability 0.5 from foreground-guided sampling targeting the rarest 50% of classes, and with probability 0.5 from repeat-factor sampling. Combining precise foreground targeting with broader crop coverage.

#### Loss Function Ablation

Under standard loss formulations, gradients are dominated by large classes, causing rare classes to contribute relatively little to optimization unless explicitly reweighted or emphasized. Loss functions that bias toward recall are therefore better suited here. We evaluate 14 configurations formed from different combinations of the loss functions as listed in this section. All experiments use repeat-factor sampling, no masking, 2D UNet, and 60 epochs with 2,000 iterations per epoch.

#### Notation

*p*_*i*_ ∈ [0, 1] is the predicted probability for pixel *i*, and *g*_*i*_ ∈ {0, 1} is the ground truth. All summations are computed over annotated pixels within a batch.

- **Dice**. Measures overlap between prediction and ground truth as 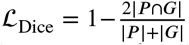. Dice loss emphasizes region-level overlap and reduces the influence of large background regions compared to pixel-wise losses Milletari et al. (2016).
- **Binary Cross Entropy (BCE) and Focal**. BCE is a pixel-wise loss that optimizes each pixel independently. Focal loss (Lin et al., 2017) extends BCE by down-weighting well-classified examples using a focusing parameter *γ*, increasing the contribution of harder examples:

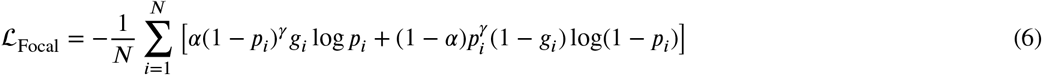

where *α* = 0.25 and *γ* = 2.0. Both BCE and Focal loss are also combined with overlap-based losses in additive formulations such as Dice+BCE, Dice+Focal, and Tversky+Focal.
- **Tversky**. Generalizes Dice by weighting false positives and false negatives separately Salehi et al. (2017):

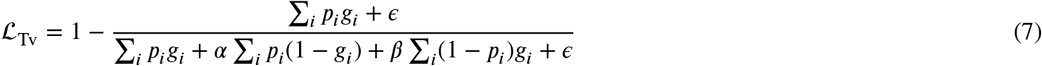

where *∈* = 10^−6^. Setting *α* < *β* penalizes false negatives more and biases the model toward recall Abraham and Khan (2019).
- **Tversky-BCE (TvBCE)**. Tversky loss is useful for imbalanced segmentation because it directly optimizes overlap while allowing different penalties for false positives and false negatives. However, for rare classes with very limited foreground overlap, a purely overlap-based objective may provide a weak training signal during the early stages of optimization. We therefore add BCE to provide dense pixel-wise supervision, while retaining the class-imbalance control of Tversky. We evaluate the same (*α, β*) settings after adding the BCE term.
- **Per-class routing**. We divide classes into common and rare groups, with each group optimized using a different loss function. The 14 common classes use Dice, while the 18 rare classes use TvBCE (*α* = 0.3, *β* = 0.7). This tests whether a frequency-aware mixed objective can outperform a single global loss.

#### Masking Strategy Ablation

In the CellMap dataset, each crop annotates only a subset of the 32 classes, and unannotated pixels are treated as NaN rather than background. Replacing these NaNs with zero incorrectly treats unannotated classes as false negatives, suppressing rare-class predictions Petit, Thome, Charnoz, Hostettler and Soler (2018). In addition, ultra-sparse classes may contain fewer than a 100 pixels within a 256 × 256 patch, making the spatial distribution of the gradient signal critical. Masking strategies address both issues by controlling which pixels contribute to the loss, without changing the model, data, or optimizer. All masking experiments use repeat-factor sampling, TvBCE (*α* = 0.3, *β* = 0.7), a 2D UNet, and 60 epochs with 2,000 iterations per epoch.

- **Entropy masking**.

Prior work uses prediction uncertainty to select training samples, but with different assumptions. OHEM Shrivastava, Gupta and Girshick (2016) focuses on high-loss regions, while entropy-based filtering methods Ghazal et al. (2025) discard uncertain pseudo-labels. MetaDCSeg Mu, Chen, Yang, Yang and Deng (2025) instead increases attention on ambiguous boundary pixels. A fundamental assumption of these methods is that the hardest or most uncertain regions or samples are the most useful for training. However, in vEM segmentation, we found that treating all high-uncertainty pixels as equally useful was not effective. Pixel-level uncertainty can arise for several reasons. Some uncertainty comes from sparse classes such as mitochondrial ribosomes `(mito_ribo)` and vesicles. Other uncertainty arises from classes that are difficult to resolve at FIB-SEM resolution, such as `mt_in` and nuclear pore regions. In some cases, the underlying signal is ambiguous. For example, ER exit site boundaries are often not clearly defined. These regions provide weak or unreliable supervision, so forcing additional learning from them is unlikely to improve performance.

For this reason, our entropy-masking implementation uses entropy as a learning signal rather than a simple selection rule. Instead of focusing on the hardest pixels, we prioritize regions of intermediate uncertainty where the model is actively improving. We up-weight these pixels by 1.5× and exclude only the highest-entropy pixels.

We compute per-pixel binary entropy

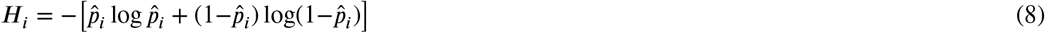

and average it across class channels. Two Exponentially Moving Average (EMA)-smoothed thresholds (τ_low_, τ_high_) divide annotated pixels into three groups, confident (*w*_*i*_ = 1.0), informative (*w*_*i*_ = 1.5), and highly uncertain (*w*_*i*_ = 0), respectively.

Fig 5 illustrates the entropy-guided curriculum. During training, we compute entropy over annotated pixels and track the 30th and 90th percentiles using an EMA with momentum 0.95. These thresholds adaptively partition pixels into low, mid, and high-entropy supervision regions. As the model becomes more confident, the boosted region progressively concentrates around evolving organelle uncertainty, while highly uncertain regions are excluded from supervision. The first 5 epochs use uniform weighting because entropy is initially high across nearly all pixels.

**Fig. 5:**
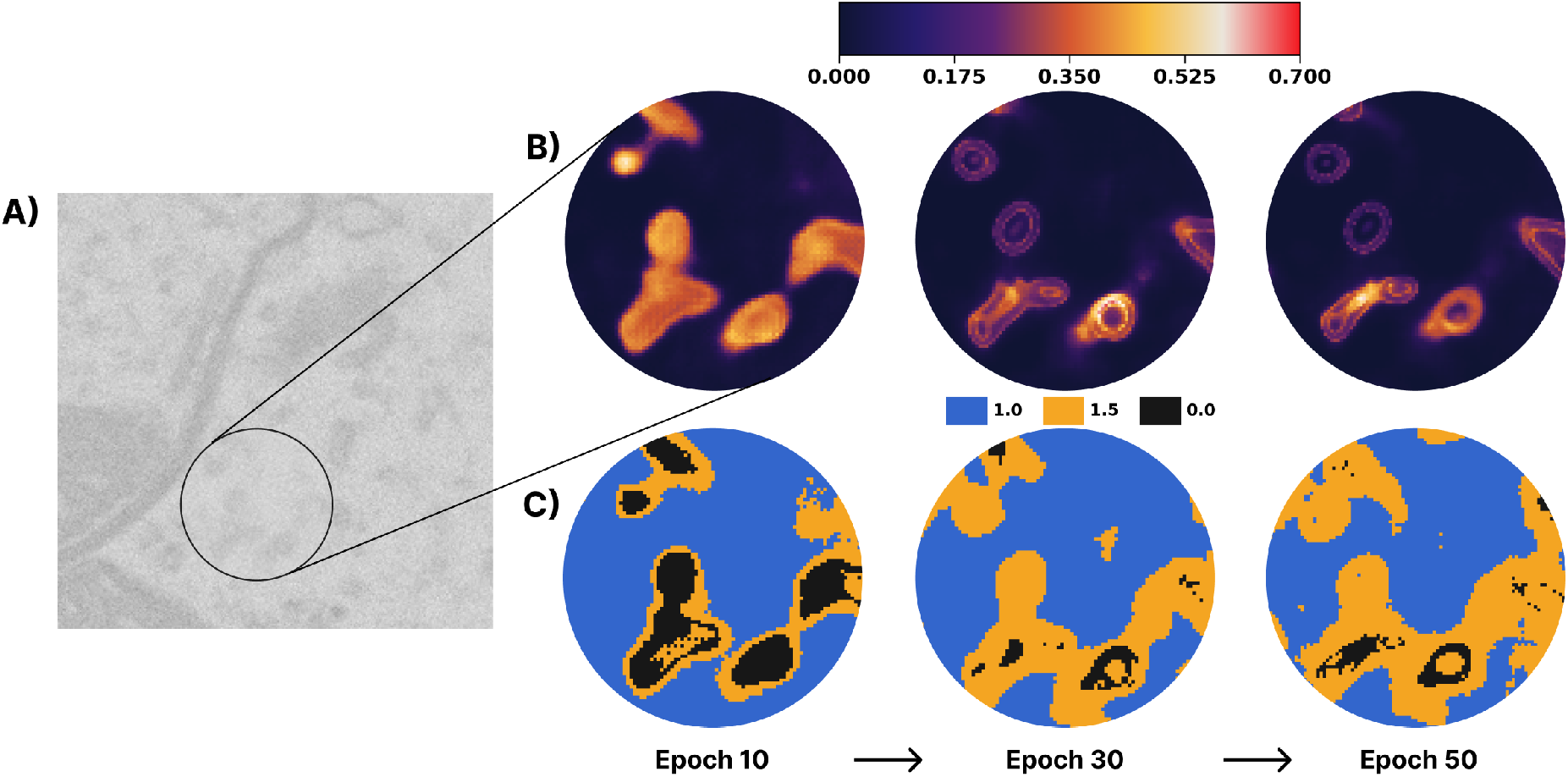
Entropy-guided masking curriculum for a zoomed ambiguous FIB-SEM region. (A) Raw input slice with the selected region of interest. (B) Entropy maps at epochs 10 and 30 show how uncertainty becomes concentrated near organelle boundaries. (C) The corresponding curriculum masks convert entropy into pixel-wise supervision weights.

- **Entropy linear-weighting**. To evaluate whether uncertain regions should be emphasized rather than excluded, we implement entropy linear-weighting. This strategy keeps all annotated pixels in the loss while assigning larger weights to pixels with higher prediction uncertainty. We compute the per-pixel binary entropy *H*_*i*_ as above and normalize it to [0, 1] within each batch, yielding 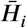. We then assign each annotated pixel the following weight

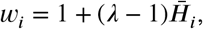

with *λ* = 2.0. This assigns weights near 1.0 to confident pixels and weights up to 2.0 to maximally uncertain pixels. Unlike entropy exclusion, which removes uncertain pixels from supervision, entropy linear-weighting preserves supervision everywhere while increasing the gradient contribution from uncertain regions. A 5-epoch warmup is applied before enabling entropy-based weighting.
- **Entropy exclusion**. To isolate the effect of removing highly uncertain pixels, we implement entropy exclusion as a simplified version of entropy masking. This strategy excludes the top 10% highest-entropy annotated pixels from supervision, but does not increase the weight of mid-entropy pixels and does not use EMA-smoothed thresholds. We compute the per-pixel binary entropy *H*_*i*_ as in the previous strategies and define a batch-specific 90th percentile threshold τ_0.9_. Each annotated pixel is then assigned a binary weight,

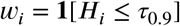

so pixels below the threshold are retained and pixels above it are excluded from the loss. The threshold is recomputed independently for each batch. A 5-epoch warmup is applied before enabling entropy-based exclusion.
- **MaskSup**. At each training step, we randomly withhold 30% of annotated pixels per channel from the main loss and used as a reconstruction target Zunair and Hamza (2022). The total loss is *L* = *L*_vis_(**w**_vis_) + 0.5 *L*_rec_(**w**_rec_), where **w**_vis_ and **w**_rec_ are complementary masks over annotated pixels. Unannotated pixels are excluded from both branches. The reconstruction branch encourages the model to infer missing pixels from surrounding context, acting as a regularizer.
- **Bounding-box masking**. Inspired by box-driven class-wise masking Song et al. (2019), we compute a tight axis-aligned bounding box around the positively annotated pixels for each class channel and pad it by 10% of its size. We assign pixels inside the box a weight of *w*_*i*_ = 1.0 and pixels outside the box a weight of *w*_*i*_ = 0.05. This concentrates the gradient near foreground regions. However, it assumes objects are small and localized. For large classes such as cyto and ecs, the box spans most of the image, so the masking has little effect. As a result, the method is inconsistent across classes and provides limited overall benefit.

#### Architecture and Pretraining Ablation

In the architecture ablation, we keep the training recipe fixed and vary only the model configuration. We use repeat-factor sampling, TvBCE (*α*=0.3, *β*=0.7), entropy masking, and train for 60 epochs with 4,000 iterations per epoch. The batch size is 16 with a gradient accumulation factor of 4.

We compare five configurations across encoder type and initialization:

- **UNet-scratch**: A standard 2-D UNet with DoubleConv encoder blocks (64 → 128 → 256 → 512 → 1024 channels, ∼31 M parameters), trained from scratch.
- **ResNet-UNet**: A UNet-style decoder with a timm-implemented ResNet-50 encoder Wightman (2019); He, Zhang, Ren and Sun (2016) (64 → 256 → 512 → 1024 → 2048 channels, ∼85.9 M parameters total), evaluated with both ImageNet-pretrained weights Russakovsky, Deng, Su, Krause, Satheesh, Ma, Huang, Karpathy, Khosla, Bernstein et al. (2015) and random initialization.
- **Swin-scratch**: Swin-V2 Base with a UNet decoder (∼102.5 M parameters), trained from scratch.
- **Swin-pretrained**: The same Swin-V2 Base architecture with ImageNet-pretrained encoder weights.

The UNet and ResNet configurations differ in both encoder structure and initialization. Only the Swin pair isolates the effect of pretraining while keeping the architecture fixed. For pretrained models, the early encoder layers are frozen. For ResNet-50, this includes `conv1, bn1,` and `layer1.` For Swin-V2, this includes the patch embedding and transformer stages 0–1. The remaining encoder layers use progressively smaller learning rates toward the input, scaled by *γ* = 0.75 per stage. The decoder, segmentation head, and input projection use the full base learning rate.

## 4. Results and Inference

Despite the extreme class imbalance in the dataset, voxel count alone does not fully explain where segmentation performance breaks down. Classes can fail due to sparsity, visual similarity to nearby structures, or geometric ambiguity at FIB-SEM resolution. Vesicle membranes, for example, appear in 178 of 214 training crops but remain difficult to segment, with the best Dice score around 0.23. This likely reflects the visual similarity between vesicular, endosomal, and lysosomal membrane classes at FIB-SEM resolution.

Understanding these failures requires looking beyond voxel frequency alone. One such factor is crop frequency. It is the number of training crops in which a class appears, and is often a better predictor of failure than total voxel count. Classes that appear in very few crops are effectively absent from most training batches regardless of their overall voxel frequency, as seen for Golgi membrane and lumen, which are each annotated in only 19 crops. Apart from crop frequency, segmentation difficulty is also influenced by morphology. Classes range from large volumetric regions to thin membranes, where even small prediction errors produce large penalties. Other classes include elongated tubules, small punctate objects, and diffuse regions that are difficult to localize consistently. In addition, some classes are inherently difficult to distinguish at FIB-SEM resolution, such as Vesicle, endosome, and lysosomal membranes which share similar visual appearance, leading to low precision. ER exit sites are small ER-associated regions whose defining structural features are often not clearly resolved, making them difficult to detect reliably.

Fig 6 shows how each ablation axis progressively improves rare-class segmentation. The baseline model trained with uniform sampling often misses sparse classes entirely, as seen for vesicles and peroxisomes. However, Repeat-factor sampling improves rare-object recall, while TvBCE further improves segmentation of thin boundaries and small classes. We also find that Entropy masking produces cleaner boundaries and reduces spurious predictions in difficult small classes such as mitochondrial ribosomes. For microtubules, lumen `(mt_in)` prediction appears only after TvBCE and entropy masking are introduced, demonstrating the cumulative effect of the training pipeline on difficult, rare classes. The following sub-sections analyze the contribution of each training axis in detail.

**Fig. 6:**
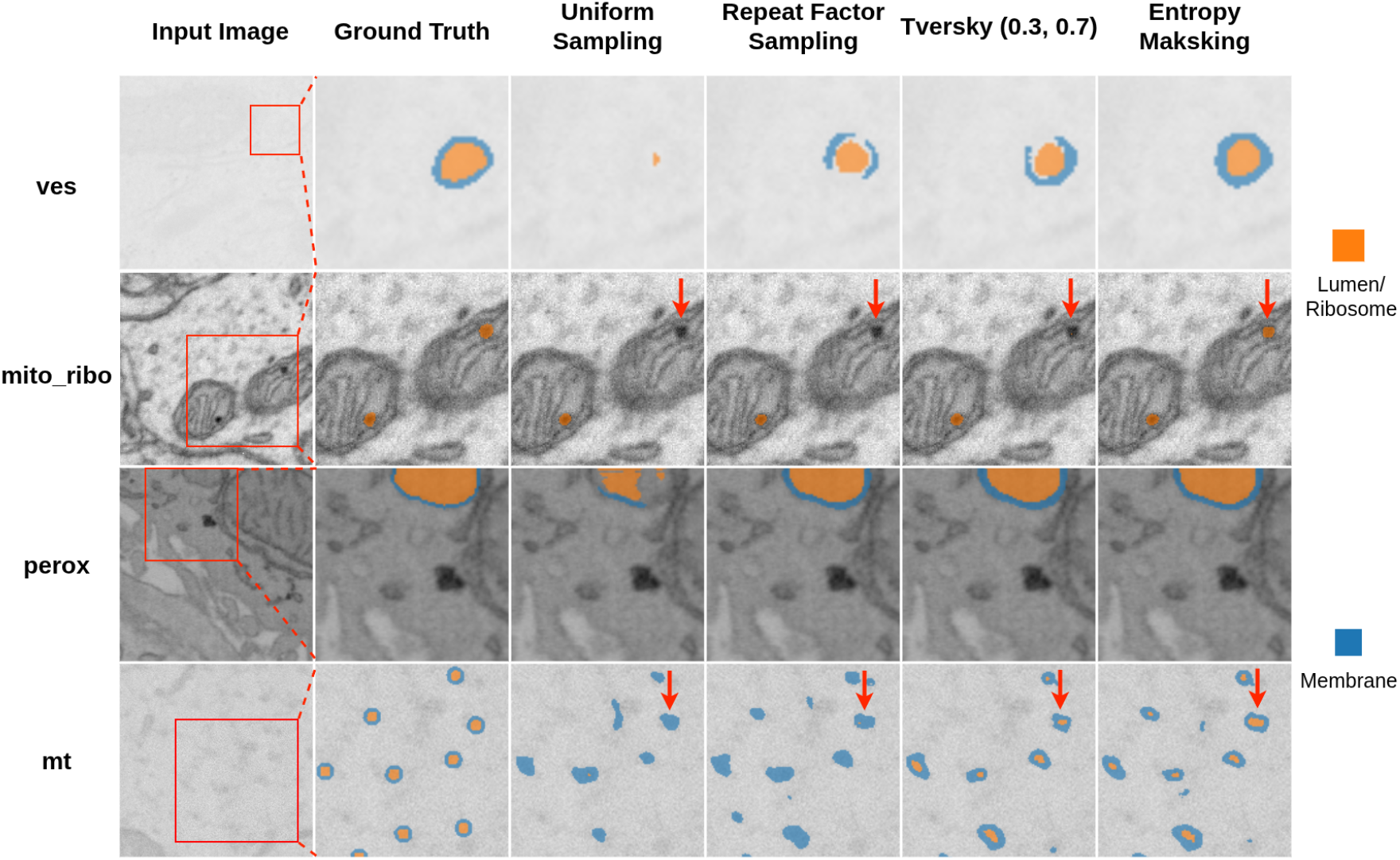
Qualitative comparison across the sequential training axes of the proposed pipeline. Each row highlights a different challenging organelle class: vesicles `(ves)`, mitochondrial ribosomes `(mito_ribo),` peroxisomes `(perox)`, and microtubules `(mt).` The input image column shows the original FIB-SEM crop with the selected region of interest. In contrast, subsequent columns show ground truth, the baseline model trained with uniform sampling, and the best-performing configuration after each successive training axis, repeat-factor sampling, TvBCE loss (*α*=0.3, *β*=0.7), and entropy masking. The progression illustrates cumulative improvements in rare-class recovery, boundary refinement, and separation of visually ambiguous classes across the training pipeline.

### 4.1 Context-Preserving Crop Sampling Outperforms Foreground-Centered Patch Mining

We find that strategies that use a centroid-based approach (foreground-guided patch mining (0.3722) and hybrid (0.3870)) to extract specific patches from the data to target rare classes during training performed the worst. These approaches reduce model performance and rank below the uniform sampling baseline (0.4960) in Dice score. In contrast, strategies that focus solely on altering crop selection probabilities outperform uniform sampling. Most of the improvement is from the rare classes. Repeat-factor sampling tops the list with a mean rare Dice of 0.3288. An important outcome here is that this difference between centroid-based patch mining approaches and crop-level sampling strategies suggests that forcing patches to center on rare foreground regions reduces contextual diversity, which is essential for multi-class segmentation.

Foreground-guided sampling performs worst among all strategies, despite appearing well-suited for rare classes in large volumetric datasets. It produces the lowest rare-class recall (0.1581), nearly half that of repeat-factor sampling (0.3653). However, its precision is not the lowest, suggesting that the main failure is reduced detection coverage rather than poor prediction quality. Models trained on centroid-centered patches fail to recognize rare classes outside the limited spatial contexts seen during training. Similar behavior is observed in the hybrid case, indicating that aggressively targeting rare foreground regions can reduce the contextual diversity required for robust multi-class segmentation.

Repeat-factor sampling improved rare-class performance relative to the other sampling strategies, yielding the highest rare-class Dice (0.3288) and rare-class precision (0.3768) in this comparison. Uniform sampling gives the highest rare-class recall (0.3735) and the best common-class Dice (0.7166), while repeat-factor sampling gives the highest common-class recall (0.7401). Overall, repeat-factor sampling provides the best balance between rare and common-class performance, achieving the highest mean Dice across all classes (0.4977), as shown in Table 2.

**Table 2.**
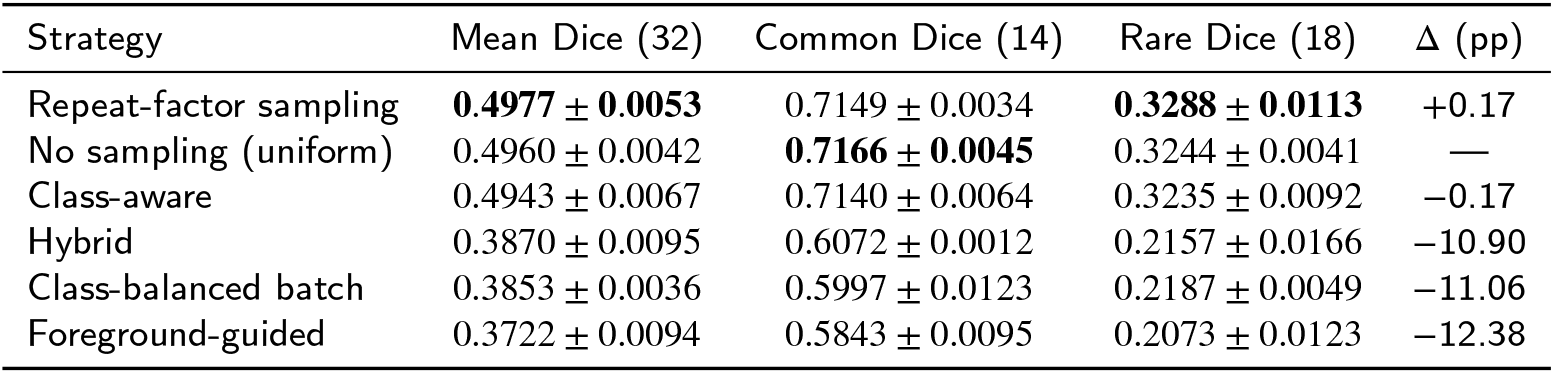
Sampling strategy comparison. Mean Dice across all 32 classes, common-class mean Dice (14 classes), and rare-class mean Dice on the validation set. Results are mean ± std over three seeds.

We find that class-level difficulty is influenced more by annotation coverage and morphology than by voxel frequency alone. Small classes that appear in many training crops and have clear visual structure (e.g., `mito_ribo)` can be learned reliably, while rare or visually ambiguous classes remain difficult despite targeted sampling. A consistent performance gap between membranes and their corresponding lumens suggests that sampling improves exposure to rare structures but does not fully resolve geometry-sensitive errors in thin boundaries. This motivates the next ablation, where we evaluate whether loss functions with stronger rare-class sensitive supervision can further improve segmentation performance.

### 4.2 Recall-Biased Loss Design Improves Rare-Class accuracy Without Sacrificing Common Classes

Table 3 reports the loss-function comparison across the 10 configurations with complete multi-seed results, while the full single-seed comparison across all 14 loss configurations is provided in Supplemental Table 7. Overall performance varies within a narrow range, from 0.4754 to 0.5007 mean Dice. Common-class Dice remains relatively stable across losses (0.7009–0.7179), while rare-class Dice shows greater sensitivity to the loss formulation (0.2999– 0.3332).

**Table 3.** Loss function comparison. Mean Dice, common-class mean Dice, and rare-class mean Dice on the validation set. All runs use repeat-factor sampling, 2D UNet, and no masking.

| Loss | Mean Dice (32) | Common Dice (14) | Rare Dice (18) |
| --- | --- | --- | --- |
| TvBCE($\alpha=0.3$, $\beta=0.7$) | **0.5007 ± 0.0166** | **0.7180 ± 0.0055** | 0.3318 ± 0.0293 |
| Tversky($\alpha=0.3$, $\beta=0.7$) | 0.4972 ± 0.0037 | 0.7080 ± 0.0062 | **0.3332 ± 0.0092** |
| TvBCE($\alpha=0.4$, $\beta=0.6$) | 0.4970 ± 0.0060 | 0.7147 ± 0.0048 | 0.3277 ± 0.0072 |
| Per-class: Dice (14 common) + TvBCE (18 rare) | 0.4970 ± 0.0088 | 0.7115 ± 0.0053 | 0.3302 ± 0.0140 |
| Dice+BCE | 0.4959 ± 0.0043 | 0.7171 ± 0.0012 | 0.3239 ± 0.0083 |
| Dice | 0.4945 ± 0.0039 | 0.7178 ± 0.0018 | 0.3209 ± 0.0074 |
| Tversky($\alpha=0.4$, $\beta=0.6$) | 0.4934 ± 0.0071 | 0.7150 ± 0.0011 | 0.3211 ± 0.0121 |
| Dice+Focal | 0.4883 ± 0.0092 | 0.7077 ± 0.0101 | 0.3177 ± 0.0099 |
| TvBCE($\alpha=0.6$, $\beta=0.4$) | 0.4849 ± 0.0145 | 0.7144 ± 0.0039 | 0.3063 ± 0.0235 |
| Tversky($\alpha=0.7$, $\beta=0.3$) | 0.4754 ± 0.0066 | 0.7009 ± 0.0039 | 0.2999 ± 0.0096 |

TvBCE with *α* = 0.3 and *β* = 0.7 achieves the best overall performance (0.5007), while Tversky loss with *α* =0.3 *β* = 0.7 gives the highest rare-class Dice (0.3332). PC-DTV (0.4970) and TvBCE with *α* = 0.4 and *β* = 0.6 (0..4970) perform similarly, indicating that several Tversky-based losses are competitive. Adding BCE supervision to Tversky with *α* = 0.3 and *β* = 0.7 slightly increases overall mean Dice from 0.4972 to 0.5007, mainly through improved common-class performance, while rare-class Dice remains comparable (0.3332 vs. 0.3318). Among the complete TvBCE variants, the recall-focused setting (*β* = 0.7) outperforms the (*β* = 0.4 setting) by 1.58 percentage points in overall mean Dice and 2.55 percentage points in rare-class Dice. Focal loss with *α* = 0.25 and *γ* = 2.0 provides limited supervision for extremely sparse classes. The focusing mechanism alone does not fully compensate for this imbalance, leading to reduced performance in rare classes.

The loss ablation shows that the choice of objective matters most for rare classes that are missed because their foreground signal is small (i.e., small classes), but still visually separable. With standard Dice loss, small membrane-bounded structures are often under-recovered because their positive voxels contribute little to the aggregate objective. Recall-biased TvBCE(*α* = 0.3, *β* = 0.7) addresses this by penalizing false negatives more strongly, leading to clear gains for vesicle classes (0.1863 → 0.2498, +0.0635) and mitochondrial boundary classes (0.2747 → 0.3275, +0.0528). Nuclear pore classes also improve, but the gain is smaller (0.5526 → 0.5747, +0.0221 with Tversky(*α* = 0.4, *β* = 0.6)), suggesting that these classes are already partly supported by distinctive spatial context. In contrast, ERES and euchromatin remain difficult across losses. ERES reaches only 0.0707 mean Dice even with PC-DTV, and euchromatin stays near zero across complete runs. Thus, loss design improves recoverable rare structures by increasing the influence of missed positives, but it cannot fully resolve classes whose boundaries are weak, sparse, or visually ambiguous.

Overall, TvBCE(*α*= 0 3, *β* = 0 7) provides the best trade-off across common and rare classes, but the class-level esults show that loss eweighting alone does not resolve all rare-class failures. We therefore use this loss configuration in the next ablation and check whether masking the training signal can further improve learning for uncertain or under-segmented structures.

### 4.3 Adaptive Entropy Masking Provides Robust Supervision for Rare-Class Segmentation

The masking ablation shows that pixel-level supervision has different effects depending on how aggressively the training signal is filtered. Among the three strongest complete-seed strategies, Entropy masking, MaskSup, and Entropy linear weighting achieve similar overall performance, with mean Dice between 0.5015 and 0.5063, Table 4. Their common-class Dice remains tightly clustered between 0.7121 and 0.7190, while rare-class Dice ranges from 0.3372 to 0.3455. MaskSup gives the highest rare-class Dice (0.3455), whereas Entropy linear weighting gives the highest rare-class recall. Entropy masking gives the best overall mean Dice (0.5063), but its rare-class Dice is slightly lower than MaskSup (0.3409 vs. 0.3455). This rare-class detection improvement is also visible in Supplemental Figure 3.

**Table 4.** Masking strategy comparison. Mean Dice, common-class mean Dice, and rare-class mean Dice on the validation set. All runs use TvBCE(*α*=0.3, *β*=0.7) and repeat-factor sampling.

| Strategy | Mean Dice (32) | Common Dice (14) | Rare Dice (18) | $\Delta$ (pp) |
| --- | --- | --- | --- | --- |
| Entropy masking | **0.5063 ± 0.0124** | **0.7190 ± 0.0058** | 0.3409 ± 0.0261 | +0.04 |
| MaskSup | 0.5059 ± 0.0042 | 0.7121 ± 0.0024 | **0.3455 ± 0.0072** | — |
| Entropy linear weighting | 0.5015 ± 0.0147 | 0.7129 ± 0.0024 | 0.3372 ± 0.0273 | −0.44 |
| Bounding-box masking | 0.4811 ± 0.0196 | 0.7040 ± 0.0098 | 0.3078 ± 0.0271 | −2.48 |
| Entropy exclusion | 0.4016 ± 0.0530 | 0.6184 ± 0.0348 | 0.2329 ± 0.0766 | −10.43 |

**Table 5.** Comparison of segmentation architectures. Metrics are mean ± std over 3 seeds. Dice averaged over all 32 classes, 14 common classes, and 18 rare classes. All models use repeat-factor sampling, entropy masking, and TvBCE loss (*α*=0.3, *β*=0.7).

| Architecture | Init | Params | Mean Dice (32) | Common Dice (14) | Rare Dice (18) |
| --- | --- | --- | --- | --- | --- |
| 2D UNet | Scratch | 31.0 M | **0.4857 ± 0.0045** | **0.7135 ± 0.0058** | **0.3086 ± 0.0079** |
| ResNet-UNet | Scratch | 85.9 M | 0.4623 ± 0.0062 | 0.6850 ± 0.0107 | 0.2891 ± 0.0087 |
| Swin-V2 + UNet | Scratch | 102.5 M | 0.4107 ± 0.0102 | 0.6411 ± 0.0077 | 0.2315 ± 0.0121 |
| ResNet-UNet | Pretrained | 85.9 M | 0.4659 ± 0.0152 | 0.6918 ± 0.0085 | 0.2903 ± 0.0220 |
| Swin-V2 + UNet | Pretrained | 102.5 M | 0.4490 ± 0.0142 | 0.6928 ± 0.0170 | 0.2594 ± 0.0121 |

**Table 6.** Best result per experimental axis. Mean Dice values are computed on the 38-crop validation set over 32 classes.

| Axis | Best configuration | Mean Dice |
| --- | --- | --- |
| Sampling | Repeat-factor sampling | $0.4977 \pm 0.0053$ |
| Loss | TvBCE($\alpha=0.3$, $\beta=0.7$) | $0.5007 \pm 0.0166$ |
| Masking | Entropy masking | $0.5063 \pm 0.0124$ |
| Architecture | 2D U-Net (scratch) | $0.4858 \pm 0.0045$ |
| **Combined best** | 2D U-Net + TvBCE($\alpha=0.3$, $\beta=0.7$) + entropy masking + repeat-factor | **$0.5063 \pm 0.0124$** |

When foreground voxels are scarce, masking strategies determine which regions receive the most supervision during training. Table 4 reports all five strategies under fixed TvBCE (*α*=0.3, *β*=0.7) and repeat-factor sampling.

Entropy masking achieves the best overall Dice (0.5063) by concentrating supervision on informative regions of the image. Moderately uncertain pixels are upweighted by 1.5×, while the pixels with highest uncertainty are excluded from supervision. By doing so, entropy masking emphasizes decision boundaries where rare classes are hard to separate and reduces the influence of noisy or ambiguous regions. As shown in Fig 7, EMA-based thresholds adapt throughout training to keep the focus aligned with these regions as the model becomes more confident. Compared with the best overall loss-only setting, Entropy masking improves ERES Dice from 0.0426 to 0.1372 for `eres_mem` and from 0.0553 to 0.0777 for `eres_lum`, although both classes remain among the most difficult due to their extremely small voxel fractions.

**Fig. 7:**
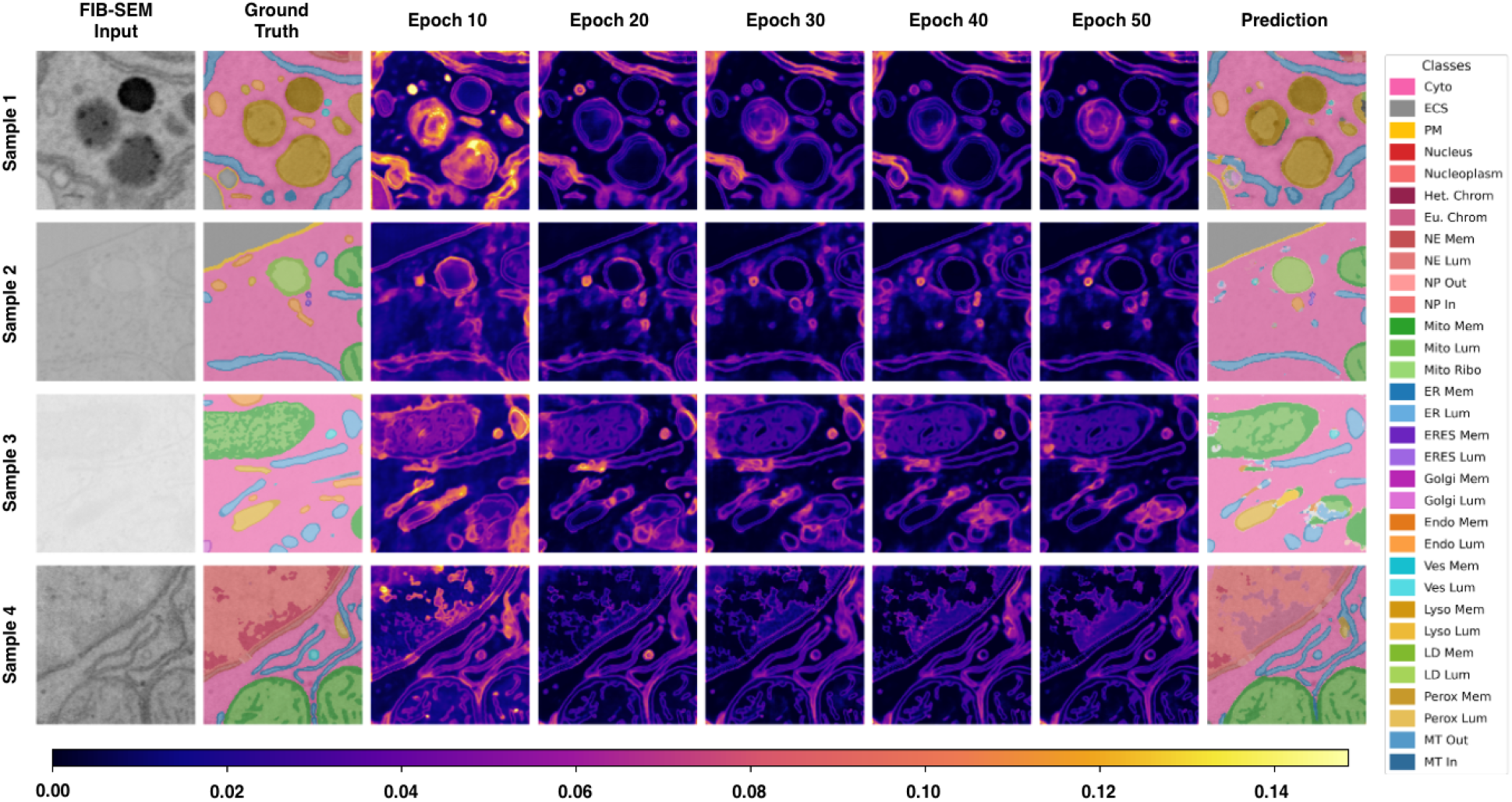
Entropy-guided masking reduces model uncertainty during training. Rows show different validation samples. Columns (left to right): raw FIB-SEM input, mean binary entropy at epochs 10–50, ground truth, and final prediction. Entropy (dark = confident, bright = uncertain) decreases substantially as training progresses, concentrating in genuinely uncertain regions by epoch 50.

Simpler entropy methods fail because they miss parts of this balance. Entropy exclusion performs substantially worse (0.4016 overall Dice, 0.2329 rare-class Dice). It removes the highest-entropy pixels but does not upweight intermediate-entropy regions, so it provides no additional supervision for uncertain but learnable boundaries. Also its thresholds are computed batch-wise rather than smoothed with EMA, the selected pixels can also vary more sharply as the entropy distribution changes during training. As a result, rare-class pixels might be excluded early, weakening their learning signal. In contrast, entropy linear weighting (0.5015) increases the contribution of all uncertain pixels, with weights scaling up to a maximum of 2.0×, increasing rare-class recall (0.4788) but lowering rare-class precision (0.3094) because noisy regions are overemphasized. Together, these results show that boosting, exclusion, and adaptive thresholds are all needed to effectively guide learning toward rare classes such as `eres_mem` and `eres_lum.`

MaskSup (0.5059) performs nearly as well as Entropy masking overall and gives the highest rare-class Dice (0..3455). By masking 30% of input pixels, it encourages the model to rely more on spatial context than local appearance, which helps context-defined structures such as the nuclear pore inner ring (`np_in`; 0.6794). Bounding-box masking wins the most per-class comparisons (10 of 32), but its overall performance is lower (0.4811). Its gains are therefore class-specific rather than broadly improving rare-class segmentation. For example, `eres_mem` reaches 0.0621 with bounding-box masking compared with 0.1372 with Entropy masking.

Masking provides targeted improvements rather than large overall gains. Compared with repeat-factor sampling, Entropy masking increases overall Dice from 0.4977 to 0.5063 and rare-class Dice from 0.3288 to 0.3409, while common-class Dice also changes only slightly from 0.7149 to 0.7190. The clearest effect is on difficult boundary classes, where pixel-level entropy can emphasize uncertain membrane regions. For example, `eres_mem` improves from 0.00 to 0.1372, compared with 0.00 to 0.0777 for `eres_lum.` Overall, entropy masking achieves the best overall Dice among masking variants (0.5063), though only marginally over MaskSup. It is also the only masking strategy that improves both common and rare-class performance relative to the best loss-only configuration.

### 4.4 Architecture and Pretraining Comparison

We compare five model configurations: `unet_scratch, resnet_scratch, resnet_pretrained, swin_scratch,` and `swin_pretrained.` Since `resnet_pretrained` uses a pretrained ResNet-50 encoder rather than the standard U-Net encoder, we interpret this experiment as a comparison of practical architecture-initialization combinations rather than a strict factorial analysis. Among the evaluated models, `unet_scratch` achieves the best overall performance with a mean Dice of 0.4858. These results also show that simply increasing the number of iterations per epoch (2000 to 4000) does not necessarily improve performance. The lower rare-class Dice in the longer architecture-ablation runs suggests that rare structures remain sensitive to training dynamics, even when common-class performance is preserved. Thus, additional training alone is not sufficient to resolve the rare-class bottleneck.

`unet_scratch` achieves the best rare-class mean Dice (0.3086) as well as the best common-class mean Dice (0. .7135). `swin_pretrained` remains competitive on common classes (0.6928), but performs substantially worse on rare classes (0.2594). This gap is driven mainly by sparse or fine-detail classes. `unet_scratch` performs better on `ld_lum, ld_mem, perox_lum,` and `perox_mem,` whereas `swin_pretrained` performs better on large, high-scoring classes such as `nuc, ecs,` and `cyto. resnet_pretrained` achieves intermediate performance (0.2903 rare-class Dice and 0.6918 common-class Dice), while `swin_scratch` performs worst on both subsets (0.2315 and 0.6411). Overall, these results suggest that unet_scratch is more effective at recovering sparse and structurally difficult classes, while is more competitive on large and visually simpler regions.

At the per-class level, the comparison is more nuanced. `unet_scratch` and `swin_pretrained` each achieve the highest Dice on 12 of 32 individual classes, but these wins occur in different parts of the label space. `swin_pretrained` performs best on several large or high-scoring structures. In contrast, `unet_scratch` performs better on several sparse or fine-detail classes. `resnet_pretrained` shows targeted strengths on lysosomal classes, achieving the best Dice for `lyso_mem` and `lyso_lum,` but these gains are not sufficient to improve its overall ranking. Taken together, the results indicate that `unet_scratch` is the strongest model on average, while different architectures show class-specific strengths. Full per-class results are provided in the Supplementary Material.

### 4.5 Summary of Findings

Across four controlled ablations, architecture, sampling, loss, and masking, the main bottleneck is consistently rare-class performance. The 18 classes with < 0.32% of training voxels drive most of the differences between configurations, while common-class Dice remains relatively stable around ∼0.71. The best pipeline combines a simple 2D U-Net trained from scratch, repeat-factor sampling (0.4977), recall-biased TvBCE loss (*α* = 0.3, *β* = 0.7; 0.5007), and entropy masking (0.5063). To our knowledge, this specific combination of strategies has not been previously applied to segmentation of volumetric EM data. The gains from each component are modest, but the strongest complete configuration is obtained when sampling, loss design, and entropy masking are combined. Per-class validation and test performance for all organelle classes is shown in Fig 8, while the corresponding numerical values are reported in Supplemental Table 2. Per-class validation Dice scores for all ablations are reported in Supplemental Tables 3–6. Qualitative comparisons are shown in Supplemental Figures 1–4, and taxonomy-level trends are summarized in Appendix C.

**Fig. 8:**
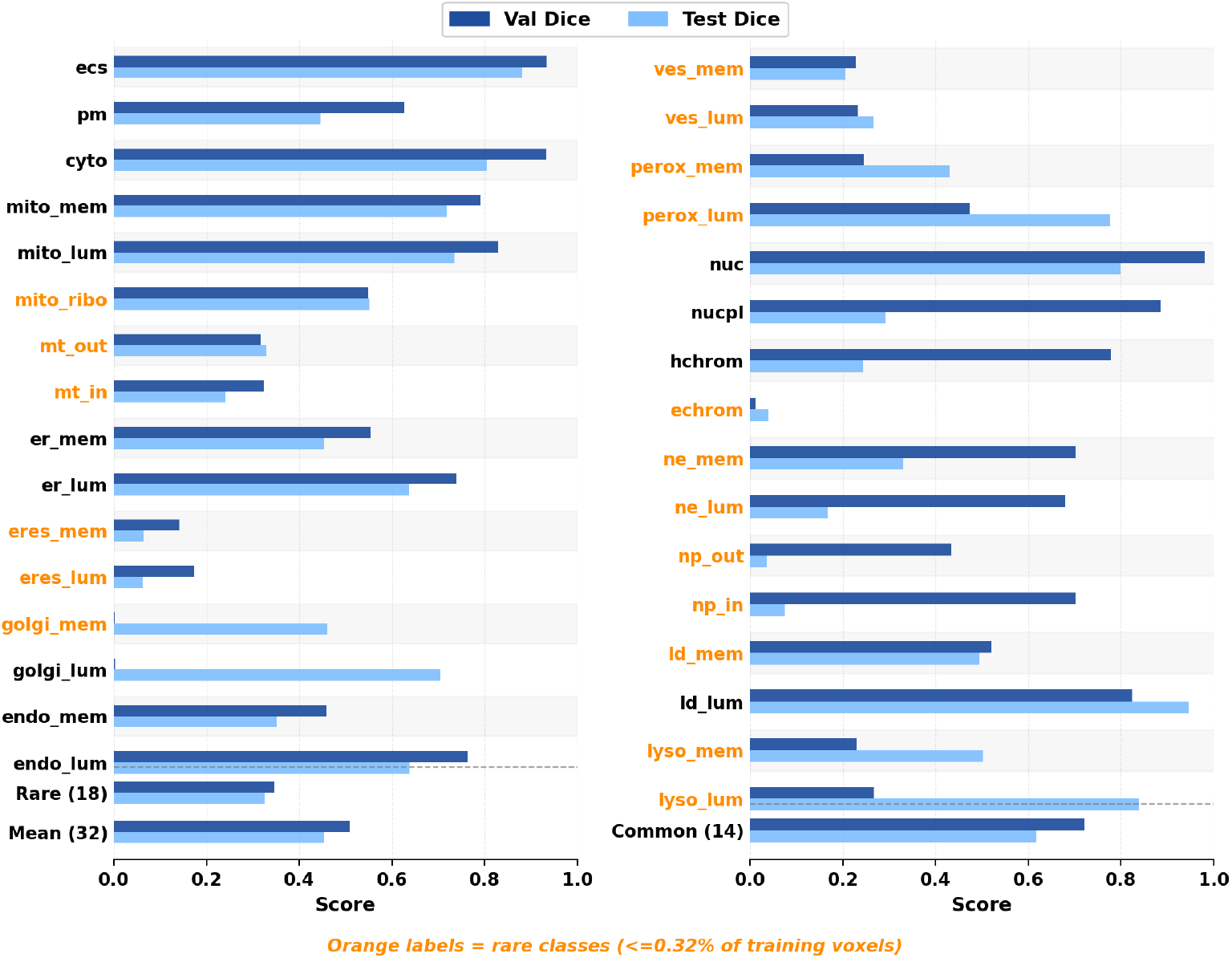
Per-class validation and test performance across all 32 organelle classes. Dice scores are shown for both splits, with rare classes (< 0.32% voxel frequency) highlighted separately.

On the held-out test set, mean Dice decreases from 0.5063 on validation to 0.4535, driven primarily by six nuclear classes: `nucpl, hchrom, ne_mem, ne_lum, np_out,` and `np_in.` These classes show the largest declines, including `np_in` (0.6749), `nucpl` (0.8814), and `hchrom` (0.7693). The remaining 26 classes show a mean change of +0.0501, but this hides mixed trends. Some classes that were near zero on validation improve sharply, such as `golgi_lum` (0.0029) and `lyso_lum` (0.2586). In contrast, common structural classes decline, including `cyto` (−0.1272), `pm` (−0.1816), and `nuc` (−0.1786), while ER exit sites remain low on both splits. This gap is consistent with split-level distribution differences. The validation set covers 38 crops from 20 volumes, while the test set contains 37 crops from 19 volumes and includes two tissue types not seen during validation. These differences may contribute to the larger decline in nuclear classes and ER exit sites, rather than indicating a uniform drop across all organelles.

Despite improvements from sampling, loss design, and entropy masking, several classes remain consistently unresolved across configurations. The most difficult failures occur in visually ambiguous classes rather than simply the rarest ones. Euchromatin `(echrom)` remains near-zero across validation and test because it is weakly separated from surrounding nucleoplasm. ER exit sites `(eres_mem, eres_lum)` also remain low, consistent with their small size and visual similarity to the ER. Vesicle, endosome, and lysosome membranes remain challenging because of their thin boundary geometry and similarity to nearby membrane structures at FIB-SEM resolution. These failures suggest that voxel frequency alone does not determine segmentation difficulty. Instead, performance is jointly limited by annotation coverage, morphology, contextual ambiguity, and the intrinsic visual separability of the target class. Future improvements may therefore require stronger structural priors, topology-aware objectives, or higher-context volumetric modeling rather than additional reweighting or oversampling alone.

## 5. Discussion

The key challenge in segmentation of volumetric data with severe class imbalance is not the “segmentation architecture”, but determining the factors that contribute to improved learning for rare and visually ambiguous classes. Major differences between methods come from the 18 rare classes, each occupying less than 0.32% of training voxels. Repeat-factor sampling gives a modest improvement over no sampling, increasing rare-class Dice from 0.3244 to 0.3288. Recall-biased TvBCE further improves rare-class Dice to 0.3318, while Entropy masking gives the best overall Dice (0.5063) and improves rare-class Dice to 0.3409. MaskSup gives the highest rare-class Dice among the masking variants (0.3455), although its overall Dice is slightly lower than Entropy masking.

These results suggest that sampling, loss design, and masking address complementary parts of the imbalance problem. Sampling controls how often rare-class crops are seen, while the loss function controls how strongly rare positive voxels influence learning. Entropy masking acts within each patch by suppressing highly uncertain pixels and emphasizing informative mid-entropy regions, which gives targeted improvements without substantially changing common-class performance. More aggressive foreground-based sampling strategies perform worse, indicating that increasing foreground exposure alone is not sufficient and may remove useful spatial context.

The loss ablation also shows that optimizing aggregate Dice and recovering the most suppressed classes are not always identical goals. For example, PC-DTV ranks below TvBCE overall (0.4970 versus 0.5007), but improves several suppressed rare classes relative to Dice-only training, including `ves_mem` (0.1569), `mt_out` (0.2641), and `mt_in` (0.2853). This supports the broader conclusion that the hardest classes are limited not only by voxel frequency, but also by morphology, annotation coverage, contextual ambiguity, and visual separability. These results show that rare-class failure reflects both limited supervision and class-specific ambiguity.

An additional strength of this benchmark is that the training and evaluation crops span multiple voxel resolutions and imaging scales. Qualitative examples in Fig 9 show that the proposed pipeline produces consistent organelle segmentations across these scale differences, suggesting that the training recipe is not restricted to a single imaging resolution. More broadly, this resolution diversity highlights the importance of developing vEM segmentation methods that remain robust across acquisition settings and explicitly account for voxel size or scale information in future work.

**Fig. 9:**
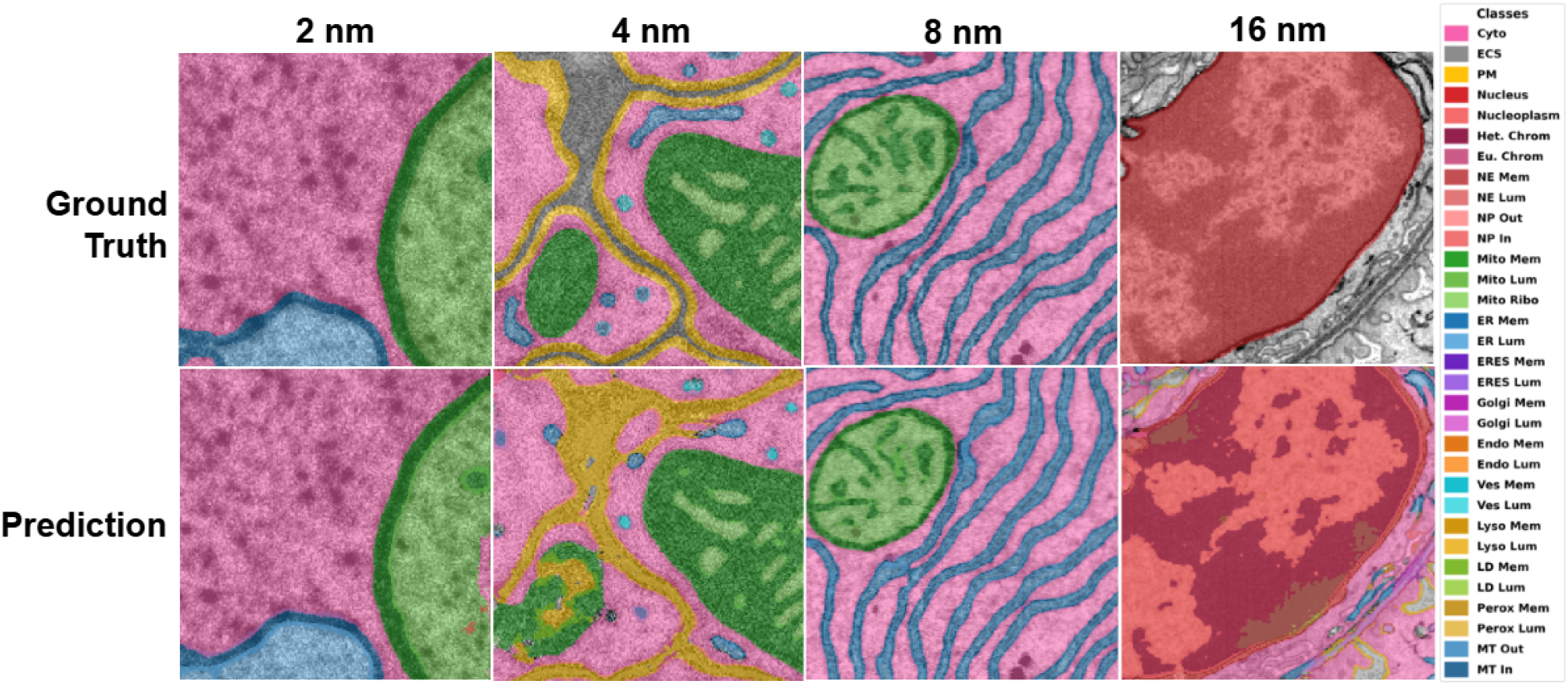
Qualitative comparison of predictions across voxel resolutions. Representative crops are shown at 2, 4, 8, and 16 nm resolution with ground-truth annotations, and model predictions displayed for each resolution.

## 6. Conclusion

This study shows that rare organelle segmentation in FIB-SEM is shaped not only by model architecture, but also by how training samples rare classes, weights errors, and selects supervised pixels. Across sampling, loss, masking, and architecture ablations, the strongest gains come from improving the training recipe. Repeat-factor sampling increases rare-class exposure, recall-biased TvBCE reduces missed detections, and entropy-guided masking refines supervision in uncertain regions. At the same time, rare-class performance cannot be explained by voxel frequency alone, as crop coverage, morphology, visual ambiguity, and class context also determine how well each class is learned. These results suggest that future studies should consider data coverage, class taxonomy, and supervision choices alongside architecture when analyzing organelle segmentation performance.

Although this study focuses on rare organelle segmentation in FIB-SEM, the same training principles may extend to other volumetric medical imaging tasks with severe class imbalance. Similar challenges arise in 3D MRI and CT segmentation of small lesions, tumors, metastases, or abnormal tissue, where clinically important regions may occupy only a small fraction of the volume. In these settings, improved rare-target sampling, recall-biased losses, and uncertainty-guided masking may help improve sensitivity without relying only on more complex architectures.

Overall, our findings support a more controlled approach to vEM segmentation benchmarks. Architecture comparisons are useful, but they are incomplete without matched sampling, loss, masking, and evaluation protocols. Future work should also focus on improving cross-tissue robustness, adding volumetric context for morphology-dependent classes, and designing supervision strategies that recover suppressed rare classes without sacrificing overall segmentation quality.

## 7. Author contributions: CRediT

**Shanmukha Vamshi Kuruba:** Conceptualization, Methodology, Software, Validation, Formal analysis, Investigation, Data curation, Visualization, Writing - original draft, Writing - review & editing. **George Stephenson:** Conceptualization, Software, Methodology, Validation, Investigation, Data curation, Writing - review. **Vignesh Kasinath:** Conceptualization, Supervision, Formal analysis, Project administration, Resources, Writing - review & editing.

## 8. Declaration of generative AI and AI-assisted technologies in the manuscript preparation process

During the preparation of this work, the authors used Grammarly in order to improve readability and grammar. We used ChatGPT for assistance with manuscript organization to comply with Medical Image Journal format. However, all aspects of the manuscript were inspected, reviewed, and edited manually by the authors. The authors take full responsibility for the content of the published article.

## 9. Declaration of Competing Interest

The authors declare that they have no known competing financial interests or personal relationships that could have appeared to influence the work reported in this paper.

## 10. Data and code availability

The data used in this study are available through the CellMap Segmentation Challenge GitHub repository. The code used for model training, evaluation, and analysis will be made publicly available upon publication.

## 11. Funding

This work was supported by funding from the National Institutes of Health, including an R35 MIRA award from the National Institute of General Medical Sciences to Vignesh Kasinath, abbreviated here as V.K., award number R35GM155426, the National Science Foundation award NSF-MCB 2446197, CU Boulder start-up funds awarded to V.K., and the Pew Charitable Trusts through V.K.’s appointment as a Pew Scholar in the Biomedical Sciences.

## Notes

https://github.com/janelia-cellmap/cellmap-segmentation-challenge

### Competing Interest Statement

The authors have declared no competing interest.

## References

Abraham, N., Khan, N.M., 2019. A novel focal tversky loss function with improved attention u-net for lesion segmentation, in: 2019 IEEE 16th international symposium on biomedical imaging (ISBI 2019), IEEE. pp. 683–687.

Brügger, R., Baumgartner, C.F., Konukoglu, E., 2019. A partially reversible u-net for memory-efficient volumetric image segmentation, in: International conference on medical image computing and computer-assisted intervention, Springer. pp. 429–437.

Buhmann, J., Sheridan, A., Malin-Mayor, C., Schlegel, P., Gerhard, S., Kazimiers, T., Krause, R., Nguyen, T.M., Heinrich, L., Lee, W.C.A., et al., 2021. Automatic detection of synaptic partners in a whole-brain drosophila electron microscopy data set. Nature methods 18, 771–774.

CellMap Consortium, 2024. Cellmap segmentation challenge dataset. doi:10.25378/janelia.c.7456966.

Ciresan, D., Giusti, A., Gambardella, L., Schmidhuber, J., 2012. Deep neural networks segment neuronal membranes in electron microscopy images. Advances in neural information processing systems 25.

Conrad, R., Narayan, K., 2023. Instance segmentation of mitochondria in electron microscopy images with a generalist deep learning model trained on a diverse dataset. Cell Systems 14, 58–71.

Ghazal, M.T., Tanha, J., Shahi, N., Roshan, S., 2025. Uncertainty-weighted semi-supervised learning with dynamic entropy masking and bhattacharyya-regularized loss. Scientific Reports .

Gupta, A., Dollar, P., Girshick, R., 2019. Lvis: A dataset for large vocabulary instance segmentation, in: Proceedings of the IEEE/CVF conference on computer vision and pattern recognition, pp. 5356–5364.

Hatamizadeh, A., Tang, Y., Nath, V., Yang, D., Myronenko, A., Landman, B., Roth, H.R., Xu, D., 2022. Unetr: Transformers for 3d medical image segmentation, in: Proceedings of the IEEE/CVF winter conference on applications of computer vision, pp. 574–584.

He, J., Zhou, G., Zhou, S., Chen, Y., 2021. Online hard patch mining using shape models and bandit algorithm for multi-organ segmentation. Ieee Journal of Biomedical and Health Informatics 26, 2648–2659.

He, K., Zhang, X., Ren, S., Sun, J., 2016. Deep residual learning for image recognition, in: Proceedings of the IEEE conference on computer vision and pattern recognition, pp. 770–778.

Heinrich, L., Bennett, D., Ackerman, D., Park, W., Bogovic, J., Eckstein, N., Petruncio, A., Clements, J., Pang, S., Xu, C.S., et al., 2021. Whole-cell organelle segmentation in volume electron microscopy. Nature 599, 141–146.

Isensee, F., Jaeger, P.F., Kohl, S.A., Petersen, J., Maier-Hein, K.H., 2021. nnu-net: a self-configuring method for deep learning-based biomedical image segmentation. Nature methods 18, 203–211.

Lin, T.Y., Goyal, P., Girshick, R., He, K., Dollár, P., 2017. Focal loss for dense object detection, in: Proceedings of the IEEE international conference on computer vision, pp. 2980–2988.

Loshchilov, I., Hutter, F., 2019. Decoupled weight decay regularization, in: International Conference on Learning Representations. URL:https://openreview.net/forum?id=Bkg6RiCqY7.

Maier-Hein, L., Reinke, A., Godau, P., Tizabi, M.D., Buettner, F., Christodoulou, E., Glocker, B., Isensee, F., Kleesiek, J., Kozubek, M., et al., 2024.Metrics reloaded: recommendations for image analysis validation. Nature methods 21, 195–212.

Milletari, F., Navab, N., Ahmadi, S.A., 2016. V-net: Fully convolutional neural networks for volumetric medical image segmentation, in: 2016 fourth international conference on 3D vision (3DV), Ieee. pp. 565–571.

Mu, C., Chen, G., Yang, X., Yang, E., Deng, C., 2025. Metadcseg: Robust medical image segmentation via meta dynamic center weighting. arXiv preprint arXiv:2511.18894 .

Peddie, C.J., Genoud, C., Kreshuk, A., Meechan, K., Micheva, K.D., Narayan, K., Pape, C., Parton, R.G., Schieber, N.L., Schwab, Y., et al., 2022. Volume electron microscopy. Nature Reviews Methods Primers 2, 51.

Petit, O., Thome, N., Charnoz, A., Hostettler, A., Soler, L., 2018. Handling missing annotations for semantic segmentation with deep convnets, in: International Workshop on Deep Learning in Medical Image Analysis, Springer. pp. 20–28.

Ronneberger, O., Fischer, P., Brox, T., 2015. U-net: Convolutional networks for biomedical image segmentation, in: International Conference on Medical image computing and computer-assisted intervention, Springer. pp. 234–241.

Russakovsky, O., Deng, J., Su, H., Krause, J., Satheesh, S., Ma, S., Huang, Z., Karpathy, A., Khosla, A., Bernstein, M., et al., 2015. Imagenet large scale visual recognition challenge. International journal of computer vision 115, 211–252.

Salehi, S.S.M., Erdogmus, D., Gholipour, A., 2017. Tversky loss function for image segmentation using 3d fully convolutional deep networks, in: International workshop on machine learning in medical imaging, Springer. pp. 379–387.

Shen, L., Lin, Z., Huang, Q., 2016. Relay backpropagation for effective learning of deep convolutional neural networks, in: European conference on computer vision, Springer. pp. 467–482.

Shrivastava, A., Gupta, A., Girshick, R., 2016. Training region-based object detectors with online hard example mining, in: Proceedings of the IEEE conference on computer vision and pattern recognition, pp. 761–769.

Smith, L.N., Topin, N., 2019. Super-convergence: Very fast training of neural networks using large learning rates, in: Artificial intelligence and machine learning for multi-domain operations applications, SPIE. pp. 369–386.

Song, C., Huang, Y., Ouyang, W., Wang, L., 2019. Box-driven class-wise region masking and filling rate guided loss for weakly supervised semantic segmentation, in: Proceedings of the IEEE/CVF Conference on Computer Vision and Pattern Recognition, pp. 3136–3145.

Sreekumar, S.P., Palanisamy, R., Swaminathan, R., 2023. Semantic segmentation of cell painted organelles using deeplabv3plus model, in: 2023 45th Annual International Conference of the IEEE Engineering in Medicine & Biology Society (EMBC), IEEE. pp. 1–4.

Wei, D., Lin, Z., Franco-Barranco, D., Wendt, N., Liu, X., Yin, W., Huang, X., Gupta, A., Jang, W.D., Wang, X., et al., 2020. Mitoem dataset: Large-scale 3d mitochondria instance segmentation from em images, in: International Conference on Medical Image Computing and Computer-Assisted Intervention, Springer. pp. 66–76.

Wightman, R., 2019. Pytorch image models. https://github.com/rwightman/pytorch-image-models. doi:10.5281/zenodo.4414861.

Xu, C.S., Hayworth, K.J., Lu, Z., Grob, P., Hassan, A.M., García-Cerdán, J.G., Niyogi, K.K., Nogales, E., Weinberg, R.J., Hess, H.F., 2017. Enhanced fib-sem systems for large-volume 3d imaging. elife 6, e25916.

Zhang, X., Lin, Z., Wang, L., Chu, Y.S., Yang, Y., Xiao, X., Lin, Y., Liu, Q., 2024. Swincell: a transformer-based framework for dense 3d cellular segmentation. bioRxiv, 2024–04.

Zhang, Y., Liao, Q., Ding, L., Zhang, J., 2022. Bridging 2d and 3d segmentation networks for computation-efficient volumetric medical image segmentation: An empirical study of 2.5 d solutions. Computerized Medical Imaging and Graphics 99, 102088.

Zhou, Z., Siddiquee, M.M.R., Tajbakhsh, N., Liang, J., 2019. Unet++: Redesigning skip connections to exploit multiscale features in image segmentation. IEEE transactions on medical imaging 39, 1856–1867.

Zunair, H., Hamza, A.B., 2022. Masked supervised learning for semantic segmentation. arXiv preprint arXiv:2210.00923.

