## Supplemental Information for "MISO: A Controlled Ablation of Masking, Initialization, Sampling, and Optimization for Segmentation in Volumetric Electron Microscopy"

***Appendix A:*** *Crop split used for training and evaluation.*

**Supplemental Table 1:** Train/validation/test split used to train and evaluate all ablation configurations.

| *Dataset* | *Train (#)* | *Train Crop number* | *Val (#)* | *Val Crops* | *Test (#)* | *Test Crops* |
|---|---|---|---|---|---|---|
| *jrc_cos7-1a* | 8 | 234, 236, 237, 239, 243, 254, 256, 257 | 2 | 247, 252 | 2 | 248, 292 |
| *jrc_cos7-1b* | 9 | 235, 238, 240, 241, 242, 245, 258, 259, 291 | 1 | 249 | 1 | 255 |
| *jrc_ctl-id8-1* | 3 | 117, 119, 130 | 1 | 118 | 1 | 116 |
| *jrc_fly-mb-1a* | 4 | 120, 121, 122, 134 | 1 | 178 | 1 | 123 |
| *jrc_fly-vnc-1* | 4 | 173, 174, 185, 79 | 1 | 78 | 1 | 176 |
| *jrc_hela-2* | 20 | 1, 113, 13, 14, 15, 155, 19, 23, 3, 4, 55, 56, 57, 58, 59, 6, 7, 8, 9, 96 | 3 | 28, 94, 95 | 3 | 16, 18, 54 |
| *jrc_hela-3* | 15 | 100, 101, 102, 111, 181, 27, 33, 34, 50, 51, 60, 62, 63, 85, 87 | 2 | 64, 65 | 2 | 61, 86 |
| *jrc_jurkat-1* | 16 | 107, 112, 126, 180, 35, 36, 37, 38, 43, 47, 66, 67, 68, 69, 91, 93 | 2 | 182, 70 | 2 | 71, 92 |
| *jrc_macrophage-2* | 14 | 110, 31, 32, 39, 42, 49, 72, 73, 74, 75, 76, 88, 89, 90 | 2 | 109, 48 | 2 | 40, 77 |
| *jrc_mus-heart-1* | 1 | 423 | 1 | 452 | 0 | — |
| *jrc_mus-kidney* | 17 | 129, 140, 141, 146, 147, 149, 156, 158, 159, 162, 163, 165, 166, 179, 221, 229, 231 | 3 | 160, 184, 230 | 3 | 148, 161, 164 |
| *jrc_mus-kidney-3* | 0 | — | 1 | 472 | 0 | — |
| *jrc_mus-kidney-glomerulus-2* | 0 | — | 1 | 421 | 0 | — |
| *jrc_mus-liver* | 18 | 131, 132, 133, 135, 136, 138, 139, 142, 143, 145, 150, 151, | 3 | 125, 144, 416 | 3 | 124, 137, 171 |

| | | | | | | |
|---|---|---|---|---|---|---|
| | | 157, 172, 175, 177, 183, 417 | | | | |
| *jrc_mus-liver-3* | 0 | — | 0 | — | 1 | 473 |
| *jrc_mus-liver-zon-1* | 32 | 267, 269, 270, 272, 273, 274, 275, 276, 277, 278, 279, 280, 282, 289, 313, 320, 322, 325, 326, 336, 337, 345, 347, 348, 349, 351, 377, 386, 407, 410, 411, 413 | 5 | 266, 268, 321, 323, 346 | 5 | 298, 319, 324, 329, 412 |
| *jrc_mus-liver-zon-2* | 15 | 333, 340, 341, 342, 353, 354, 355, 356, 357, 358, 362, 366, 369, 376, 408 | 2 | 367, 387 | 2 | 368, 370 |
| *jrc_mus-nacc-1* | 0 | — | 0 | — | 1 | 115 |
| *jrc_sum159-1* | 8 | 22, 25, 80, 81, 83, 97, 98, 99 | 2 | 20, 84 | 2 | 26, 82 |
| *jrc_sum159-4* | 14 | 187, 188, 189, 202, 203, 206, 208, 209, 210, 212, 216, 217, 218, 219 | 2 | 201, 213 | 2 | 186, 211 |
| *jrc_ut21-1413-003* | 14 | 190, 191, 192, 193, 196, 197, 198, 199, 200, 214, 224, 225, 227, 228 | 2 | 220, 222 | 2 | 195, 226 |
| *jrc_zf-cardiac-1* | 2 | 378, 379 | 1 | 380 | 1 | 381 |
| ***TOTAL*** | **214** | | **38** | | **37** | |

***Appendix B:*** *Evaluation results per class for each ablation and final validation and test scores per class.*

**Supplemental Table 2:** Per-class validation and test performance across all 32 organelle classes. Dice and IoU scores are reported for each class (Rare classes, defined as <0.32% of training voxels, are marked with †).

| ***Class*** | ***Val Dice*** | ***Test Dice*** | ***Val IoU*** | ***Test IoU*** |
|---|---|---|---|---|
| *ecs* | 0.932 ± 0.002 | 0.880 ± 0.015 | 0.873 ± 0.004 | 0.786 ± 0.024 |
| *pm* | 0.627 ± 0.001 | 0.445 ± 0.019 | 0.456 ± 0.001 | 0.286 ± 0.015 |
| *cyto* | 0.932 ± 0.001 | 0.805 ± 0.006 | 0.872 ± 0.001 | 0.673 ± 0.008 |
| *mito_mem* | 0.792 ± 0.003 | 0.718 ± 0.002 | 0.656 ± 0.004 | 0.560 ± 0.002 |
| *mito_lum* | 0.836 ± 0.002 | 0.735 ± 0.031 | 0.718 ± 0.003 | 0.582 ± 0.039 |
| *mito_ribo †* | 0.579 ± 0.081 | 0.551 ± 0.028 | 0.410 ± 0.078 | 0.381 ± 0.027 |
| *mt_out †* | 0.315 ± 0.014 | 0.329 ± 0.021 | 0.187 ± 0.010 | 0.197 ± 0.015 |
| *mt_in †* | 0.314 ± 0.023 | 0.241 ± 0.016 | 0.186 ± 0.016 | 0.137 ± 0.010 |
| *er_mem* | 0.554 ± 0.009 | 0.452 ± 0.003 | 0.383 ± 0.008 | 0.292 ± 0.002 |

| | | | | |
|---|---|---|---|---|
| *er_lum* | 0.738 ± 0.004 | 0.637 ± 0.009 | 0.585 ± 0.005 | 0.467 ± 0.010 |
| *eres_mem †* | 0.137 ± 0.072 | 0.065 ± 0.092 | 0.075 ± 0.043 | 0.035 ± 0.050 |
| *eres_lum †* | 0.078 ± 0.052 | 0.062 ± 0.057 | 0.041 ± 0.029 | 0.033 ± 0.030 |
| *golgi_mem †* | 0.001 ± 0.001 | 0.460 ± 0.014 | 0.001 ± 0.001 | 0.299 ± 0.012 |
| *golgi_lum* | 0.003 ± 0.002 | 0.705 ± 0.014 | 0.001 ± 0.001 | 0.544 ± 0.017 |
| *endo_mem* | 0.465 ± 0.018 | 0.351 ± 0.058 | 0.303 ± 0.015 | 0.214 ± 0.043 |
| *endo_lum* | 0.762 ± 0.003 | 0.638 ± 0.016 | 0.615 ± 0.003 | 0.469 ± 0.017 |
| *ves_mem †* | 0.237 ± 0.021 | 0.204 ± 0.013 | 0.135 ± 0.013 | 0.114 ± 0.008 |
| *ves_lum †* | 0.240 ± 0.010 | 0.265 ± 0.011 | 0.136 ± 0.006 | 0.153 ± 0.007 |
| *perox_mem †* | 0.260 ± 0.177 | 0.430 ± 0.008 | 0.158 ± 0.125 | 0.274 ± 0.006 |
| *perox_lum †* | 0.470 ± 0.171 | 0.777 ± 0.012 | 0.318 ± 0.157 | 0.635 ± 0.017 |
| *nuc* | 0.978 ± 0.005 | 0.799 ± 0.004 | 0.956 ± 0.009 | 0.665 ± 0.005 |
| *nucpl* | 0.881 ± 0.003 | 0.292 ± 0.001 | 0.788 ± 0.005 | 0.171 ± 0.000 |
| *hchrom* | 0.769 ± 0.005 | 0.244 ± 0.003 | 0.625 ± 0.007 | 0.139 ± 0.002 |
| *echrom †* | 0.011 ± 0.002 | 0.039 ± 0.008 | 0.006 ± 0.001 | 0.020 ± 0.004 |
| *ne_mem †* | 0.699 ± 0.008 | 0.330 ± 0.013 | 0.537 ± 0.009 | 0.197 ± 0.009 |
| *ne_lum †* | 0.673 ± 0.011 | 0.167 ± 0.011 | 0.507 ± 0.013 | 0.091 ± 0.006 |
| *np_out †* | 0.438 ± 0.007 | 0.036 ± 0.010 | 0.280 ± 0.006 | 0.019 ± 0.005 |
| *np_in †* | 0.675 ± 0.009 | 0.075 ± 0.009 | 0.509 ± 0.010 | 0.039 ± 0.005 |
| *ld_mem †* | 0.520 ± 0.025 | 0.495 ± 0.012 | 0.351 ± 0.023 | 0.329 ± 0.011 |
| *ld_lum* | 0.797 ± 0.065 | 0.945 ± 0.011 | 0.666 ± 0.089 | 0.897 ± 0.019 |
| *lyso_mem †* | 0.231 ± 0.018 | 0.502 ± 0.016 | 0.131 ± 0.012 | 0.335 ± 0.014 |
| *lyso_lum †* | 0.259 ± 0.006 | 0.838 ± 0.024 | 0.149 ± 0.004 | 0.722 ± 0.035 |

**Supplemental Table 3:** Per class Dice scores for individual configurations in Sampling Ablation.

| ***Class*** | ***repeat_factor*** | ***baseline*** | ***no_sampling*** | ***class_balanced*** | ***hybrid*** | ***foreground_guided*** |
|---|---|---|---|---|---|---|
| *ecs* | **0.9315** | 0.9187 | 0.9200 | 0.8249 | 0.8040 | 0.7701 |
| *pm* | 0.6176 | 0.6064 | **0.6184** | 0.5444 | 0.4986 | 0.5192 |
| *cyto* | **0.9192** | 0.9152 | 0.9183 | 0.8643 | 0.8184 | 0.8553 |
| *mito_mem* | 0.7722 | 0.7617 | **0.7762** | 0.6840 | 0.5747 | 0.7005 |
| *mito_lum* | 0.7998 | 0.7980 | **0.8229** | 0.7419 | 0.5471 | 0.7393 |
| ***mito_ribo*** | 0.6412 | **0.6581** | 0.6531 | 0.3185 | 0.3756 | 0.2991 |

| | | | | | | |
|---|---|---|---|---|---|---|
| ***mt_out*** | **0.3224** | 0.3089 | 0.2992 | 0.2788 | 0.2154 | 0.2449 |
| ***mt_in*** | **0.3205** | 0.3060 | 0.2688 | 0.2741 | 0.2748 | 0.2070 |
| *er_mem* | **0.5435** | 0.5332 | 0.5419 | 0.4960 | 0.4051 | 0.4760 |
| *er_lum* | 0.7123 | 0.7027 | **0.7201** | 0.6487 | 0.5101 | 0.6218 |
| ***eres_mem*** | 0.0000 | 0.0467 | **0.0784** | 0.0000 | 0.0048 | 0.0263 |
| ***eres_lum*** | 0.0000 | **0.0089** | 0.0000 | 0.0000 | 0.0000 | 0.0086 |
| ***golgi_mem*** | 0.0010 | 0.0002 | 0.0008 | **0.0030** | 0.0000 | 0.0001 |
| *golgi_lum* | 0.0020 | 0.0017 | 0.0012 | **0.0060** | 0.0005 | 0.0000 |
| *endo_mem* | 0.4644 | 0.4667 | **0.4719** | 0.3906 | 0.3531 | 0.3586 |
| *endo_lum* | 0.7579 | **0.7688** | 0.7644 | 0.6434 | 0.5768 | 0.6321 |
| ***lyso_mem*** | **0.3069** | 0.2850 | 0.2721 | 0.1741 | 0.1977 | 0.1763 |
| ***lyso_lum*** | 0.3282 | **0.3446** | 0.2792 | 0.1579 | 0.2444 | 0.1989 |
| ***ld_mem*** | 0.4408 | 0.4884 | **0.5393** | 0.4788 | 0.4320 | 0.4536 |
| *ld_lum* | 0.7880 | 0.8161 | 0.8203 | 0.6833 | **0.8356** | 0.6053 |
| ***ves_mem*** | 0.1573 | 0.1587 | 0.1617 | **0.1841** | 0.1359 | 0.1327 |
| ***ves_lum*** | 0.2212 | 0.2063 | 0.2064 | **0.2303** | 0.1950 | 0.1704 |
| ***perox_mem*** | **0.3586** | 0.2254 | 0.2000 | 0.2623 | 0.0751 | 0.0241 |
| ***perox_lum*** | 0.5692 | 0.5031 | 0.3721 | **0.5718** | 0.1827 | 0.0358 |
| *nuc* | 0.9628 | 0.9494 | **0.9657** | 0.7155 | 0.6201 | 0.6911 |
| *nucpl* | 0.8542 | 0.8387 | **0.8817** | 0.7476 | 0.7934 | 0.8323 |
| *hchrom* | **0.8157** | 0.7980 | 0.8039 | 0.5127 | 0.6871 | 0.6249 |
| ***echrom*** | 0.0147 | 0.0293 | 0.0281 | 0.0260 | **0.0867** | 0.0230 |
| ***ne_mem*** | 0.7097 | **0.7315** | 0.7088 | 0.4411 | 0.5581 | 0.4487 |
| ***ne_lum*** | 0.7107 | **0.7173** | 0.6926 | 0.5344 | 0.6334 | 0.5409 |
| ***np_out*** | 0.4399 | **0.4603** | 0.4523 | 0.3688 | 0.3750 | 0.3568 |
| ***np_in*** | 0.6588 | **0.7135** | 0.6735 | 0.5502 | 0.6298 | 0.5135 |

**Supplemental Table 4:** Per class validation Dice scores for top 10 individual configurations in Loss Function Ablation. Where Tversky-XY is alpha=X and beta=Y, and BCE is Binary Cross Entropy.

| *Class* | *Tversky-37+BCE* | *Per Class* | *Tversky-64+BCE* | *Dice + BCE* | *Tversky-46 BCE* | *Dice* | *Tversky-73* | *Tversky-37* | *Tversky-46* | *Dice + Focal* |
|---|---|---|---|---|---|---|---|---|---|---|
| *ecs* | 0.935 | 0.926 | 0.935 | 0.933 | 0.936 | 0.926 | 0.905 | 0.925 | 0.921 | 0.926 |
| *pm* | 0.624 | 0.635 | 0.621 | 0.63 | 0.629 | 0.628 | 0.596 | 0.609 | 0.616 | 0.628 |
| *cyto* | 0.93 | 0.925 | 0.931 | 0.932 | 0.933 | 0.921 | 0.905 | 0.917 | 0.917 | 0.927 |
| *mito_mem* | 0.785 | 0.781 | 0.781 | 0.787 | 0.79 | 0.766 | 0.735 | 0.772 | 0.767 | 0.766 |
| *mito_lum* | 0.818 | 0.829 | 0.826 | 0.84 | 0.838 | 0.802 | 0.786 | 0.813 | 0.805 | 0.812 |
| *mito_ribo* | 0.651 | 0.573 | 0.597 | 0.584 | 0.642 | 0.651 | 0.625 | 0.653 | 0.655 | 0.648 |
| *mt_out* | 0.337 | 0.329 | 0.27 | 0.299 | 0.314 | 0.255 | 0.309 | 0.33 | 0.307 | 0.292 |
| *mt_in* | 0.346 | 0.332 | 0.25 | 0.303 | 0.319 | 0.288 | 0.31 | 0.326 | 0.31 | 0.277 |
| *er_mem* | 0.558 | 0.552 | 0.551 | 0.557 | 0.561 | 0.549 | 0.508 | 0.53 | 0.531 | 0.553 |
| *er_lum* | 0.743 | 0.72 | 0.731 | 0.733 | 0.745 | 0.714 | 0.635 | 0.716 | 0.702 | 0.732 |
| *eres_mem* | 0.102 | 0.223 | 0.033 | 0 | 0 | 0 | 0.018 | 0 | 0 | 0 |
| *eres_lum* | 0.141 | 0.287 | 0.125 | 0 | 0 | 0 | 0.007 | 0 | 0 | 0 |
| *golgi_mem* | 0.001 | 0 | 0 | 0 | 0 | 0 | 0.003 | 0.002 | 0 | 0.002 |
| *golgi_lum* | 0.002 | 0.001 | 0.001 | 0 | 0 | 0 | 0.009 | 0.003 | 0 | 0.003 |
| *endo_mem* | 0.458 | 0.474 | 0.465 | 0.47 | 0.472 | 0.462 | 0.446 | 0.434 | 0.433 | 0.47 |
| *endo_lum* | 0.756 | 0.739 | 0.776 | 0.773 | 0.778 | 0.762 | 0.71 | 0.752 | 0.702 | 0.772 |
| *lyso_mem* | 0.255 | 0.269 | 0.216 | 0.232 | 0.238 | 0.221 | 0.243 | 0.26 | 0.328 | 0.282 |
| *lyso_lum* | 0.257 | 0.248 | 0.195 | 0.266 | 0.227 | 0.267 | 0.28 | 0.29 | 0.332 | 0.302 |
| *ld_mem* | 0.518 | 0.532 | 0.449 | 0.496 | 0.469 | 0.514 | 0.48 | 0.532 | 0.499 | 0.5 |
| *ld_lum* | 0.738 | 0.787 | 0.732 | 0.754 | 0.694 | 0.837 | 0.874 | 0.84 | 0.833 | 0.625 |
| *ves_mem* | 0.222 | 0.215 | 0.213 | 0.127 | 0.19 | 0.164 | 0.118 | 0.225 | 0.186 | 0.163 |
| *ves_lum* | 0.244 | 0.236 | 0.227 | 0.191 | 0.219 | 0.224 | 0.18 | 0.262 | 0.226 | 0.239 |
| *perox_mem* | 0.38 | 0.236 | 0.27 | 0.318 | 0.223 | 0.196 | 0.348 | 0.13 | 0.162 | 0.185 |
| *perox_lum* | 0.666 | 0.39 | 0.712 | 0.548 | 0.487 | 0.531 | 0.705 | 0.289 | 0.299 | 0.373 |
| *nuc* | 0.969 | 0.971 | 0.967 | 0.975 | 0.973 | 0.957 | 0.94 | 0.97 | 0.949 | 0.965 |
| *nucpl* | 0.875 | 0.865 | 0.86 | 0.858 | 0.871 | 0.865 | 0.819 | 0.889 | 0.848 | 0.867 |
| *hchrom* | 0.786 | 0.803 | 0.803 | 0.78 | 0.8 | 0.809 | 0.814 | 0.786 | 0.77 | 0.785 |
| *echrom* | 0.022 | 0.012 | 0.019 | 0.013 | 0.013 | 0.018 | 0.03 | 0.021 | 0.03 | 0.015 |
| *ne_mem* | 0.695 | 0.702 | 0.686 | 0.714 | 0.699 | 0.709 | 0.651 | 0.697 | 0.71 | 0.702 |
| *ne_lum* | 0.666 | 0.687 | 0.732 | 0.725 | 0.702 | 0.714 | 0.689 | 0.668 | 0.685 | 0.712 |
| *np_out* | 0.426 | 0.45 | 0.456 | 0.444 | 0.454 | 0.435 | 0.458 | 0.453 | 0.449 | 0.465 |
| *np_in* | 0.661 | 0.659 | 0.669 | 0.663 | 0.68 | 0.65 | 0.654 | 0.68 | 0.695 | 0.67 |

**Supplemental Table 5:** Per class Dice scores for individual configurations in Masking Ablation.

| *Class* | *Entropy Curriculum* | *Mask Supervision* | *Entropy linear weighting* | *Boundary box* | *Entropy exclusion* |
|---|---|---|---|---|---|
| *ecs* | 0.9334 | 0.9349 | 0.9267 | 0.9383 | 0.928 |

| | | | | | |
|---|---|---|---|---|---|
| *pm* | 0.6255 | 0.6267 | 0.6104 | 0.6181 | 0.6027 |
| *cyto* | 0.9307 | 0.9307 | 0.93 | 0.9361 | 0.9181 |
| *mito_mem* | 0.7892 | 0.7857 | 0.7873 | 0.7909 | 0.7798 |
| *mito_lum* | 0.8149 | 0.8133 | 0.8289 | 0.8216 | 0.8315 |
| *mito_ribo* | 0.4911 | 0.6267 | 0.5904 | 0.2442 | 0.638 |
| *mt_out* | 0.3331 | 0.3178 | 0.3326 | 0.3061 | 0.1899 |
| *mt_in* | 0.3446 | 0.3295 | 0.3454 | 0.2928 | 0.1461 |
| *er_mem* | 0.5621 | 0.556 | 0.5532 | 0.5737 | 0.4157 |
| *er_lum* | 0.7399 | 0.732 | 0.7342 | 0.761 | 0.6908 |
| *eres_mem* | 0.2524 | 0 | 0.1621 | 0.0598 | 0.0554 |
| *eres_lum* | 0.1597 | 0 | 0.1365 | 0.042 | 0.0447 |
| *golgi_mem* | 0 | 0.0031 | 0.002 | 0.0025 | 0.0002 |
| *golgi_lum* | 0 | 0.0054 | 0.0046 | 0.002 | 0.0022 |
| *endo_mem* | 0.4399 | 0.4451 | 0.4282 | 0.3702 | 0.4522 |
| *endo_lum* | 0.7537 | 0.7459 | 0.6853 | 0.6991 | 0.7761 |
| *lyso_mem* | 0.238 | 0.2943 | 0.2478 | 0.2792 | 0.261 |
| *lyso_lum* | 0.2474 | 0.279 | 0.2543 | 0.3622 | 0.034 |
| *ld_mem* | 0.5295 | 0.525 | 0.5328 | 0.5284 | 0.4947 |
| *ld_lum* | 0.7452 | 0.6678 | 0.838 | 0.8106 | 0.616 |
| *ves_mem* | 0.261 | 0.2559 | 0.2158 | 0.2325 | 0.1516 |
| *ves_lum* | 0.2525 | 0.2694 | 0.2535 | 0.2303 | 0.1796 |
| *perox_mem* | 0.498 | 0.4433 | 0.3406 | 0.2951 | 0.1779 |
| *perox_lum* | 0.6718 | 0.6681 | 0.6351 | 0.6486 | 0.5059 |
| *nuc* | 0.9747 | 0.9724 | 0.9778 | 0.9859 | 0.9824 |
| *nucpl* | 0.8726 | 0.8731 | 0.8909 | 0.8926 | 0.8615 |
| *hchrom* | 0.7646 | 0.7745 | 0.7606 | 0.7994 | 0.771 |
| *echrom* | 0.0111 | 0.0192 | 0.0189 | 0.038 | 0.0064 |
| *ne_mem* | 0.7031 | 0.7008 | 0.6867 | 0.6655 | 0.643 |
| *ne_lum* | 0.6876 | 0.6799 | 0.6433 | 0.6846 | 0.6922 |
| *np_out* | 0.4353 | 0.4408 | 0.422 | 0.4188 | 0.0906 |
| *np_in* | 0.6682 | 0.6915 | 0.6262 | 0.637 | 0.0628 |

**Supplemental Table 6:** Per class Dice scores for individual configurations in Architecture and Pretraining Comparison.

| ***Class*** | ***UNet Scratch*** | ***Swin-ImageNet Pretrained*** | ***ResNet-50 Pretrained*** | ***Swin Scratch*** |
|---|---|---|---|---|
| *ecs* | 0.9332 | 0.9408 | 0.9187 | 0.8992 |
| *pm* | 0.6223 | 0.6055 | 0.5909 | 0.5335 |
| *cyto* | 0.933 | 0.9377 | 0.924 | 0.9099 |
| *mito_mem* | 0.7881 | 0.7897 | 0.7427 | 0.7188 |
| *mito_lum* | 0.8326 | 0.8624 | 0.8255 | 0.7506 |

| | | | | |
|---|---|---|---|---|
| *mito_ribo* | 0.6144 | 0.4227 | 0.6567 | 0.3779 |
| *mt_out* | 0.304 | 0.2955 | 0.3006 | 0.256 |
| *mt_in* | 0.2939 | 0.2862 | 0.3097 | 0.2331 |
| *er_mem* | 0.5489 | 0.547 | 0.5215 | 0.5043 |
| *er_lum* | 0.7254 | 0.7506 | 0.7069 | 0.6641 |
| *eres_mem* | 0.0478 | 0 | 0.0496 | 0 |
| *eres_lum* | 0.0838 | 0 | 0 | 0 |
| *golgi_mem* | 0.002 | 0 | 0 | 0 |
| *golgi_lum* | 0.0055 | 0 | 0 | 0.0028 |
| *endo_mem* | 0.4444 | 0.447 | 0.4437 | 0.3528 |
| *endo_lum* | 0.7307 | 0.7431 | 0.7079 | 0.5796 |
| *lyso_mem* | 0.2693 | 0.2824 | 0.2757 | 0.2106 |
| *lyso_lum* | 0.281 | 0.3447 | 0.4536 | 0.3908 |
| *ld_mem* | 0.552 | 0.4919 | 0.298 | 0.3538 |
| *ld_lum* | 0.8228 | 0.5922 | 0.4296 | 0.484 |
| *ves_mem* | 0.2004 | 0.2371 | 0.204 | 0.139 |
| *ves_lum* | 0.2043 | 0.2215 | 0.211 | 0.1338 |
| *perox_mem* | 0.1822 | 0.0005 | 0.0839 | 0.059 |
| *perox_lum* | 0.38 | 0.1181 | 0.1569 | 0.089 |
| *nuc* | 0.9726 | 0.9983 | 0.9959 | 0.983 |
| *nucpl* | 0.9022 | 0.9228 | 0.8911 | 0.8833 |
| *hchrom* | 0.7869 | 0.8032 | 0.7302 | 0.7644 |
| *echrom* | 0.0106 | 0.0143 | 0.0975 | 0.1607 |
| *ne_mem* | 0.6767 | 0.7236 | 0.6825 | 0.6684 |
| *ne_lum* | 0.6744 | 0.7003 | 0.64 | 0.6218 |
| *np_out* | 0.4425 | 0.4041 | 0.4523 | 0.3451 |
| *np_in* | 0.6571 | 0.5973 | 0.676 | 0.583 |

**Supplemental Table 7:** Loss function comparison. Mean Dice, common-class mean Dice, and rare-class mean Dice on the validation set for a single-seed run. All configurations use repeat-factor sampling, a 2D UNet, and no masking.

| ***Loss*** | ***Mean Dice (32)*** | ***Common Dice (14)*** | ***Rare Dice (18)*** |
|---|---|---|---|
| *TvBCE (alpha=0.3, beta=0.7)* | 0.5177 | 0.7126 | 0.3661 |
| *Per-class: Dice (14 common) + TvBCE (18 rare)* | 0.5122 | 0.7149 | 0.3545 |
| *TvBCE (alpha=0.6, beta=0.4)* | 0.5031 | 0.7129 | 0.34 |
| *Dice+BCE* | 0.4983 | 0.7158 | 0.3291 |
| *TvBCE (alpha=0.4, beta=0.6)* | 0.4968 | 0.7157 | 0.3265 |
| *Dice* | 0.4948 | 0.7141 | 0.3242 |
| *Tversky (alpha=0.7, beta=0.3)* | 0.4935 | 0.6917 | 0.3394 |

| | | | |
|---|---|---|---|
| *Tversky (alpha=0.3, beta=0.7)* | 0.493 | 0.7112 | 0.3233 |
| *Tversky (alpha=0.4, beta=0.6)* | 0.4896 | 0.6995 | 0.3263 |
| *Dice+Focal* | 0.4893 | 0.7023 | 0.3236 |
| *Tversky+Focal* | 0.4876 | 0.6969 | 0.3248 |
| *TvBCE (alpha=0.7, beta=0.3)* | 0.4772 | 0.7005 | 0.3035 |
| *BCE* | 0.4198 | 0.7077 | 0.1959 |
| *Focal (gamma=2.0)* | 0.338 | 0.649 | 0.0962 |

***Appendix C:*** *Class Taxonomy and Ablation Summary*

The 32 retained classes vary widely in voxel frequency, crop coverage, morphology, and visual distinctiveness. Voxel frequency is useful for defining rare classes, but it does not fully explain segmentation difficulty. Some classes are rare by voxel count but appear in enough training crops to be learnable, while others remain difficult because they are visually ambiguous, spatially sparse, or weakly separated from nearby structures. To capture these differences, we group classes using five criteria: voxel frequency, physical size, crop frequency, morphology, and confusability. These criteria provide a more complete explanation of class-level performance than voxel frequency alone.

**Rare and common classes**

Classes are defined as rare if they occupy approximately less than 0.32\% of the training voxels. This gives 18 rare classes and 14 common classes. Common classes, such as cytoplasm, extracellular space, nucleus, mitochondria, and ER, provide abundant training signal and perform consistently well across methods. Rare classes show larger variation across ablations and largely determine the ranking of methods.

**Physical size**

Large classes include bulk structures such as cytoplasm, extracellular space, nucleus, mitochondria, ER, and nucleoplasm. These classes span broad regions and are generally easier to learn. Medium classes include plasma membrane, heterochromatin, endosome, and lipid droplet lumen. These have moderate spatial extent and intermediate difficulty. Small classes include thin membranes, nuclear pores, microtubules, vesicles, peroxisomes, ER exit sites, and euchromatin. These classes are more sensitive to localization errors and often provide limited foreground signal.

**Crop frequency**

Crop frequency captures a different source of difficulty by measuring how many training crops contain a given class. This is not always reflected by voxel frequency. For example, mitochondrial ribosomes are extremely rare by voxel count but appear in 47 of 214 training crops, making them more learnable than their voxel frequency alone would suggest. In contrast, classes such as Golgi membrane/lumen, lipid droplet membrane/lumen, peroxisome membrane/lumen, ER exit sites, and nuclear pores appear in fewer crops and receive more limited training exposure. This shows that annotation coverage strongly affects learnability.

**Morphology**

Volumetric structures are more forgiving because small boundary errors still leave many correct interior pixels. Thin membranes are harder because even a small shift can greatly reduce overlap. Tubular structures such as microtubules are difficult because they are narrow and extended. Punctate or ring-like structures such as mitochondrial ribosomes and nuclear pores can be learned when they are visually distinctive and sufficiently represented across crops. Diffuse or ambiguous structures such as euchromatin and ER exit sites remain difficult because their boundaries are weak or not clearly visible in FIB-SEM intensity.

**Confusability**

Some classes are difficult because they are visually confusable with nearby structures. Vesicle, endosome, and lysosome membranes have similar local appearance, so the model can confuse them without broader context. The nuclear envelope is continuous with the ER, making some ER and nuclear-envelope regions hard to separate locally. ER exit sites are small ER-associated regions whose defining features are not always visible at FIB-SEM resolution. Euchromatin is also difficult because its boundary with nucleoplasm is diffuse and low contrast.

**Class Taxonomy analysis for each ablation**

**Sampling Ablation**

The sampling ablation shows that increasing rare-class exposure is helpful, but not uniformly beneficial for every rare or small class. Uniform sampling remains strong for many established classes, while repeat-factor sampling provides targeted gains for several underrepresented structures, including Golgi classes, nuclear-envelope classes, microtubules, and peroxisomes. Repeat-factor sampling improves overall Dice from 0.4960 with no sampling to 0.4977, and rare-class Dice from 0.3244 to 0.3288. More aggressive strategies such as class-balanced batching, foreground-guided sampling, and hybrid sampling perform substantially worse, suggesting that rare-class exposure must be balanced against preserving spatial context.

**Supplemental Figure 1:** Qualitative comparison of sampling strategies

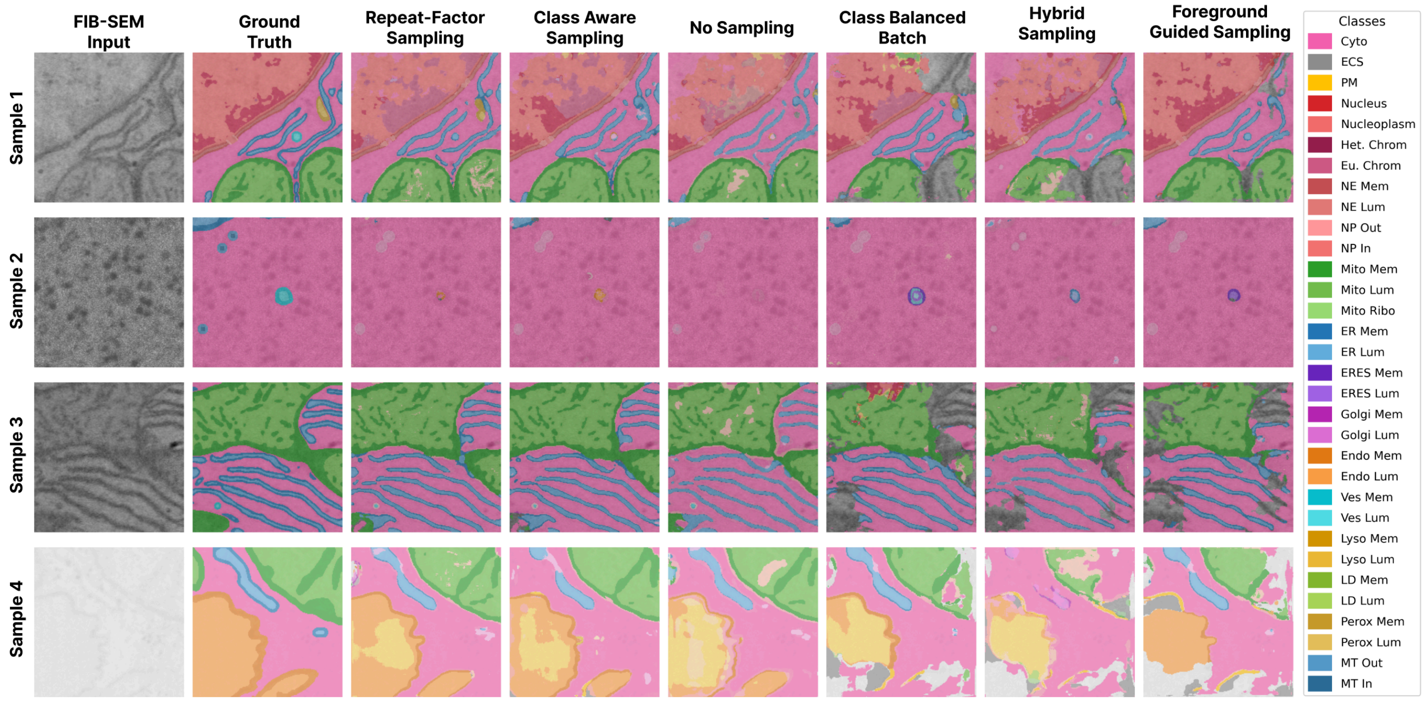


The qualitative comparison of sampling strategies in **Supplemental Figure 1** is consistent with the quantitative results. Repeat-factor sampling gives the most stable predictions among the sampling methods, while foreground-guided, class-balanced, and hybrid sampling often lose spatial context and produce weaker segmentations in dense cytoplasmic regions. Shared failures remain for isolated rare structures, showing that sampling alone cannot recover all sparse classes.

### Loss Function Ablation

The loss ablation shows that recall-biased TvBCE( alpha=0.3, beta=0.7) gives the best overall loss-ablation result (0.5007). This loss improves rare-class Dice from 0.3209 with Dice-only training to 0.3318, while common-class Dice remains essentially unchanged. PC-DTV ranks slightly lower overall (0.4970), but provides targeted gains for some suppressed rare classes, including ves_mem (0.1569 to 0.2270), mt_out (0.2641 to 0.3238), and mt_in (0.2853 to 0.3298). This shows that optimizing aggregate Dice and recovering the most suppressed rare classes are not always identical objectives.

**Supplemental Figure 2:** Qualitative comparison of the top 6 loss functions

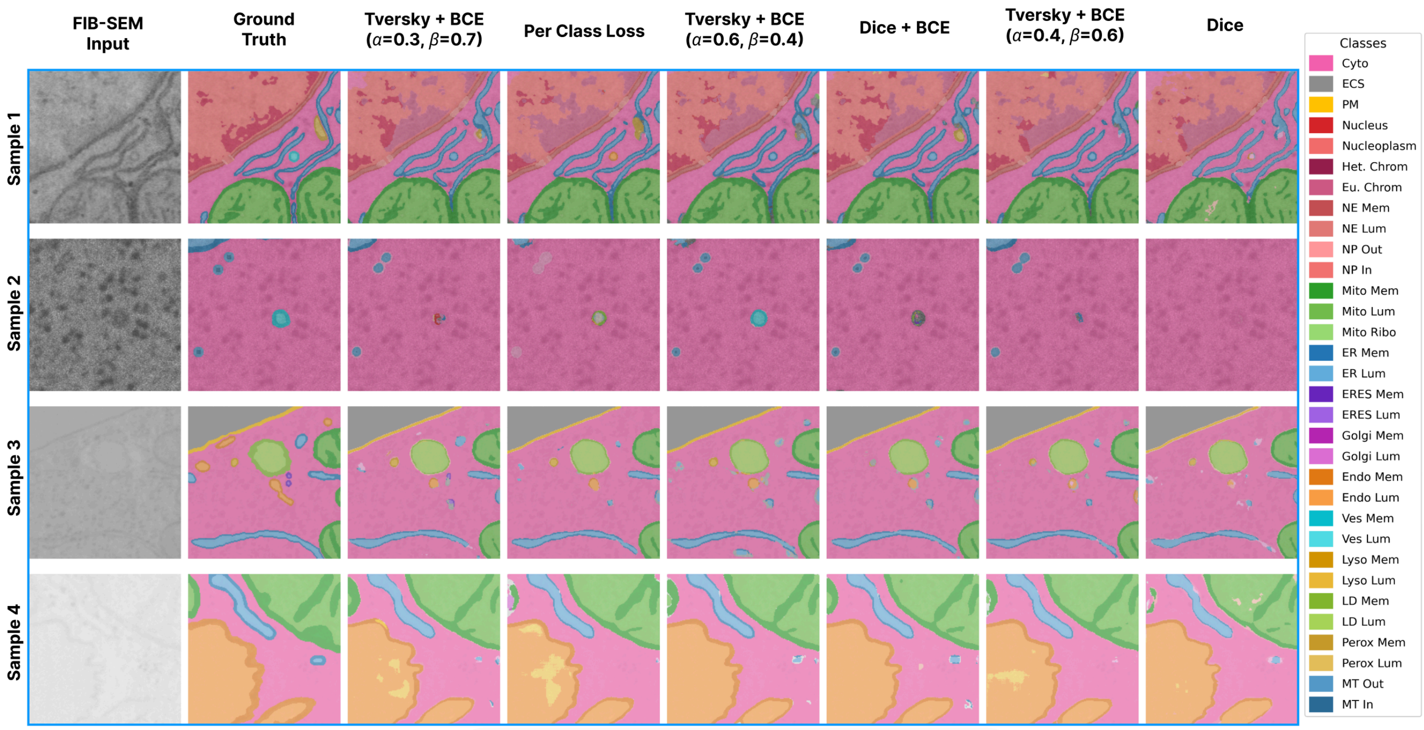


The qualitative comparison in **Supplemental Figure 2** supports this interpretation. Recall-biased and per-class losses recover more rare thin structures than plain Dice in selected examples, while plain Dice often favors larger, easier regions. However, the qualitative results also show that isolated membrane-bounded objects remain sensitive to the exact loss formulation, so loss design improves rare organelles but does not fully solve sparse-class recovery.

**Masking Ablation**

The masking ablation shows that masking methods provide targeted improvements rather than uniform gains across all classes. Entropy masking gives the best overall masking result (0.5063), while MaskSup gives the highest rare-class Dice among the masking variants (0.3455). Entropy masking performs best for mito_ribo and several membrane or boundary-associated classes, while MaskSup performs best for several context-dependent classes, including vesicle membrane/lumen and nuclear pore classes. Entropy linear weighting performs best for several lipid droplet, microtubule, and peroxisome classes. These results show that different masking strategies emphasize different structures, even though entropy masking is the best overall strategy.

**Supplemental Figure 3:** Qualitative comparison of self-supervised masking strategies

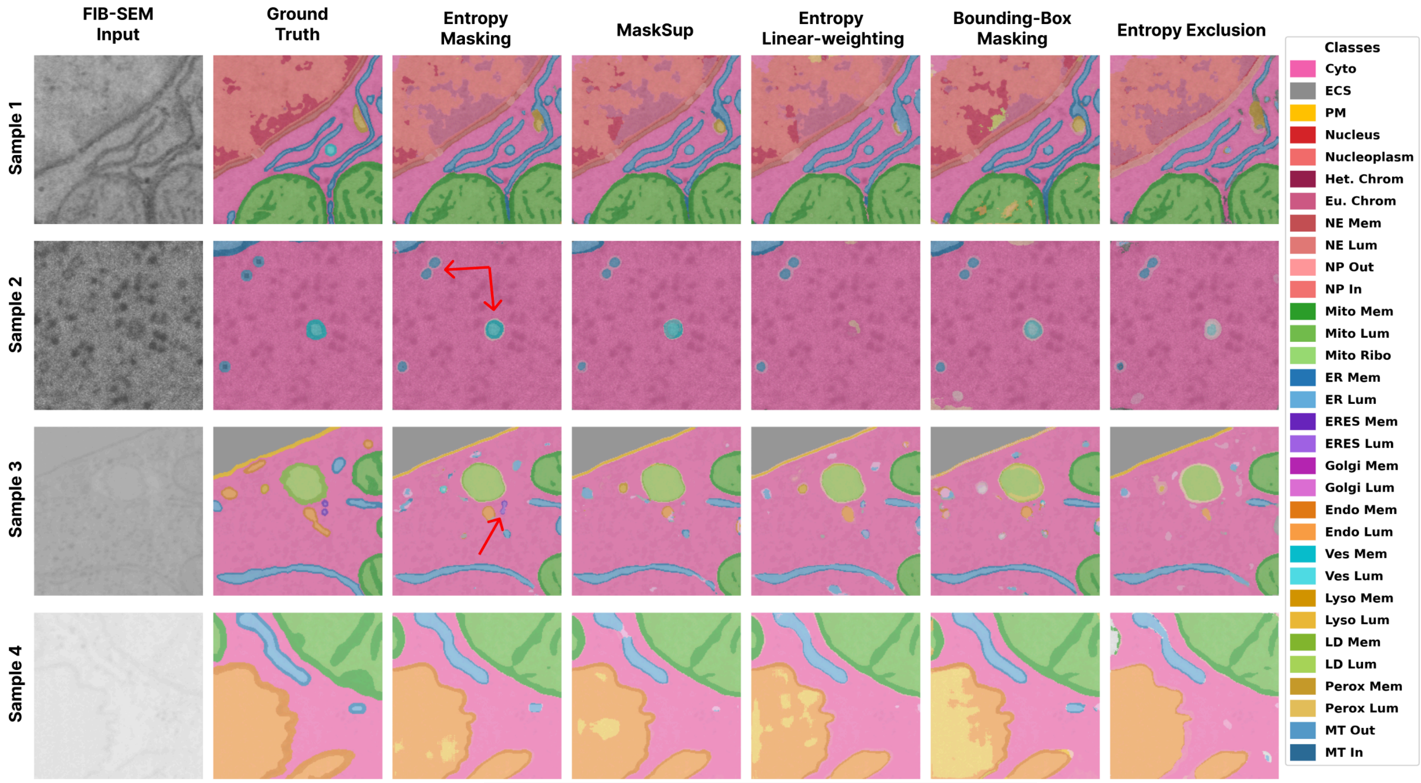


The qualitative comparison in **Supplemental Figure 3** is consistent with the class-level results. Entropy masking, MaskSup, and entropy linear weighting recover different subsets of isolated rare structures, including vesicle and microtubule classes. Entropy masking is especially useful for ambiguous boundary regions, while MaskSup is more helpful for classes that depend on local context. These examples illustrate why masking improves rare-class learning, but also why no single masking strategy dominates every class.

#### Architecture Ablation

The architecture ablation shows that larger or pretrained backbones do not automatically improve rare-class segmentation. The 2D U-Net trained from scratch achieves the best overall architecture result (0.4858), as well as the best rare-class and common-class means within the architecture comparison. At the per-class level, unet_scratch and swin_pretrained each achieve the highest Dice on 12 of 32 classes, but their strengths differ. swin_pretrained performs best on several large or high-scoring structures, including ecs, nuc, cyto, mito_mem, and mito_lum. In contrast, unet_scratch performs better on several sparse or fine-detail classes, including mito_ribo, lipid droplet classes, nuclear pore classes, and microtubules. Peroxisomes are a targeted strength of the ResNet-UNet scratch model, showing that some rare structures benefit from different architecture choices.

**Supplemental Figure 4:** Qualitative comparison of encoder architectures

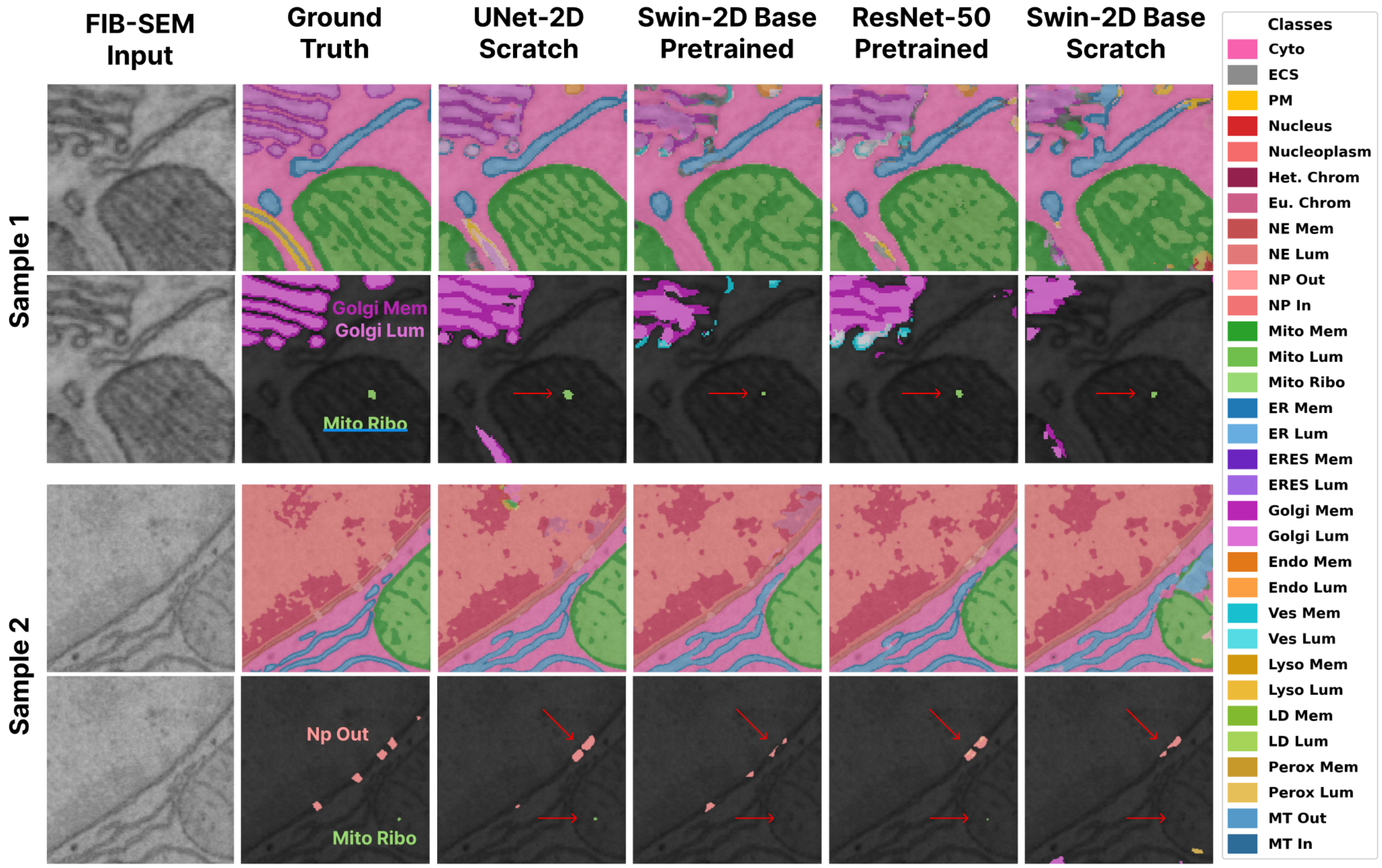


The qualitative architecture comparison in **Supplemental Figure 4** illustrates these class-specific differences. UNet-2D from scratch and pretrained Swin-V2 perform similarly on many common structures, but they differ more clearly on rare and fine-detail classes. The UNet predictions better preserve several small structures in the selected examples, while Swin-V2 remains competitive on broad, high-scoring regions. These results support the main conclusion that architecture choice matters, but rare-class recovery depends strongly on the full training recipe and class-specific structure.
